# High-resolution cryo-electron microscopy of the human CDK-activating kinase for structure-based drug design

**DOI:** 10.1101/2023.04.07.536029

**Authors:** Victoria I. Cushing, Adrian F. Koh, Junjie Feng, Kaste Jurgaityte, Ash K. Bahl, Simak Ali, Abhay Kotecha, Basil J. Greber

## Abstract

Rational design of next-generation therapeutics can be facilitated by high-resolution structures of drug targets bound to small-molecule inhibitors. However, application of structure-based methods to macromolecules refractory to crystallisation has been hampered by the often-limiting resolution and throughput of cryogenic electron microscopy (cryo-EM). Here, we use high-resolution cryo-EM to determine structures of the CDK-activating kinase, a master regulator of cell growth and division, in its free and nucleotide-bound states and in complex with 14 inhibitors at up to 1.8 Å resolution. Our structures provide detailed insight into inhibitor interactions and networks of water molecules in the active site of cyclin-dependent kinase 7. Our data support a previously proposed mechanism contributing to inhibitor selectivity, thereby providing the basis for rational design of next-generation therapeutics. Additionally, our results establish a methodological framework for the use of high-resolution cryo-EM in structure-based drug design.

## Introduction

The human CDK-activating kinase (CAK) is a heterotrimeric protein complex formed by CDK7, cyclin H, and MAT1 and acts as a master regulator of cell growth and division ^1^. It regulates transcription initiation and the cell cycle by phosphorylating RNA polymerase II and cyclin-dependent kinases, respectively. Due to this central role in cellular physiology, the CAK has been identified as a promising target for cancer therapeutics (reviewed in ref. ^2^) and it is a possible target for antivirals ^3^. Several groups have discovered specific inhibitors of the CAK ^3–,8^, including high-affinity inhibitors that occupy the active site to compete with adenosine- nucleotide binding, and covalent inhibitors that modify a cysteine residue in the vicinity of the active site. We have developed a CDK7 inhibitor series and advanced one of these compounds, ICEC0942 (CT7001, samuraciclib) ^4^, into clinical trials for treatment of advanced-stage solid cancers (clinicaltrials.gov NCT04802759, NCT03363893). To date, five other CDK7 inhibitors have entered clinical trials: SY-1365 (NCT03134638), SY-5609 (NCT04247126, NCT04929223), LY-3405105 (NCT03770494), Q901 (NCT05394103) and XL-102 (NCT04726332). One of the major challenges in the development of CDK-targeting compounds is specificity ^9^ because 20 different enzymes in human cells are classified as CDKs and often exhibit high sequence identity near their catalytic sites ^10, 11^. To enable the discovery and rational design of next-generation therapeutics with increased potency and reduced off-target effects, structural data permitting the application of structure-based drug design approaches are instrumental.

Working towards the goal of making the human CAK accessible to structure-based drug design, we previously determined its cryo-EM structure bound to nucleotide analogues and the inhibitors ICEC0942 and THZ1 ^12, 13^. Our ICEC0942-bound structure revealed interesting conformational differences between the inhibitor bound to its clinical target, CAK, and bound to CDK2, an off-target complex with the potential to cause side effects in patients, highlighting the potential of using structural data to guide the development of next-generation inhibitors in this system ^12, 14^. Our published cryo-EM structures used the human proteins and have reached higher resolution than X-ray crystal structures of a homologous complex from a thermophilic fungus ^15^, indicating that cryo-EM is the method of choice in this system, provided that throughput and resolution reach the levels required for drug discovery applications.

We therefore set out to structurally characterise complexes of CAK bound to a range of both commercially available molecules and the series of compounds developed and characterised alongside ICEC0942, aiming to uncover the structural basis of CDK7 inhibitor selectivity to pave the way towards next-generation therapeutics. The nature of this endeavour required high resolution for this small, asymmetric complex as well as high throughput, creating a technical challenge and situating this effort at the edge of what is currently feasible using cryo-EM ^16^. To address this challenge, we used rapid screening on a 200 kV cryo-transmission electron microscope (cryo-TEM) to obtain initial structures and identify promising specimens and progressed suitable specimens to high-end data collection on an energy-filtered 300 kV cryo-TEM with a cold field emission gun (cold-FEG). We have obtained 16 structures of the 85 kDa CDK-cyclin-module of the human CAK in its nucleotide-bound, free, and inhibitor bound states at up to 1.8 Å resolution. In addition to achieving high resolution from large datasets, we established routine ∼4 Å and ∼3 Å-resolution structure determination of ligand-bound complexes using the 200 kV setup combined with on-the-fly data processing from 1 hour and 4 hours of data collection. This work provides a workflow for the application of cryo-EM to structure-based drug discovery, expands our understanding of how structurally diverse inhibitors interact with the active site of CDK7, and thereby provides a basis for the design of next-generation cancer therapeutics.

## Results

### High-resolution structures of the human CAK

Structure-based drug design greatly benefits from high-resolution data that provide molecular models with atomic accuracy and information on water networks in the active sites of target complexes. Knowledge of ordered water positions is critical for understanding of inhibitor binding ^17^ and can provide avenues towards inhibitor optimisation (e.g. ^18, 19^). Our previous structure of nucleotide-bound human CAK reached 2.8 Å resolution ^13^, limiting its ability to resolve many ordered waters, and an apo-structure of the fully assembled human CAK is lacking altogether. We therefore set out to determine the high-resolution structures of apo- and nucleotide-bound human CAK using the latest generation Titan Krios cryo-TEM ^20^. Furthermore, as a proof of concept for high-resolution structure determination of inhibitor-bound CAK, we re-determined the structure of CAK modified by the covalent inhibitor THZ1 (Extended Data. Fig. 1a), which has been shown to disrupt super-enhancer transcription in human cancer cell lines ^7^, and we determined the structure of CAK in complex with the high-affinity inhibitor LDC4297 (Extended Data. Fig. 1b), a pyrazolotriazine-class compound that exhibits anti-viral activity ^3^, a property that is of particular interest given current global health challenges.

These experiments resulted in cryo-EM reconstructions of the ligand-bound human CAK at 1.9-2.1 Å resolution (Fig. 1a, Extended Data Fig. 1c-e). The maps show protein side chain and main chain features consistent with the reported resolution (Fig. 1b, c), allow direct identification of post-translational modifications, such as N-terminal protein acetylation (Fig. 1d), and reveal the locations of many water molecules (Fig. 1b-d). At high resolution, density for the regulatory CDK7 T-loop becomes fragmented, indicating structural heterogeneity.

**Figure 1.**
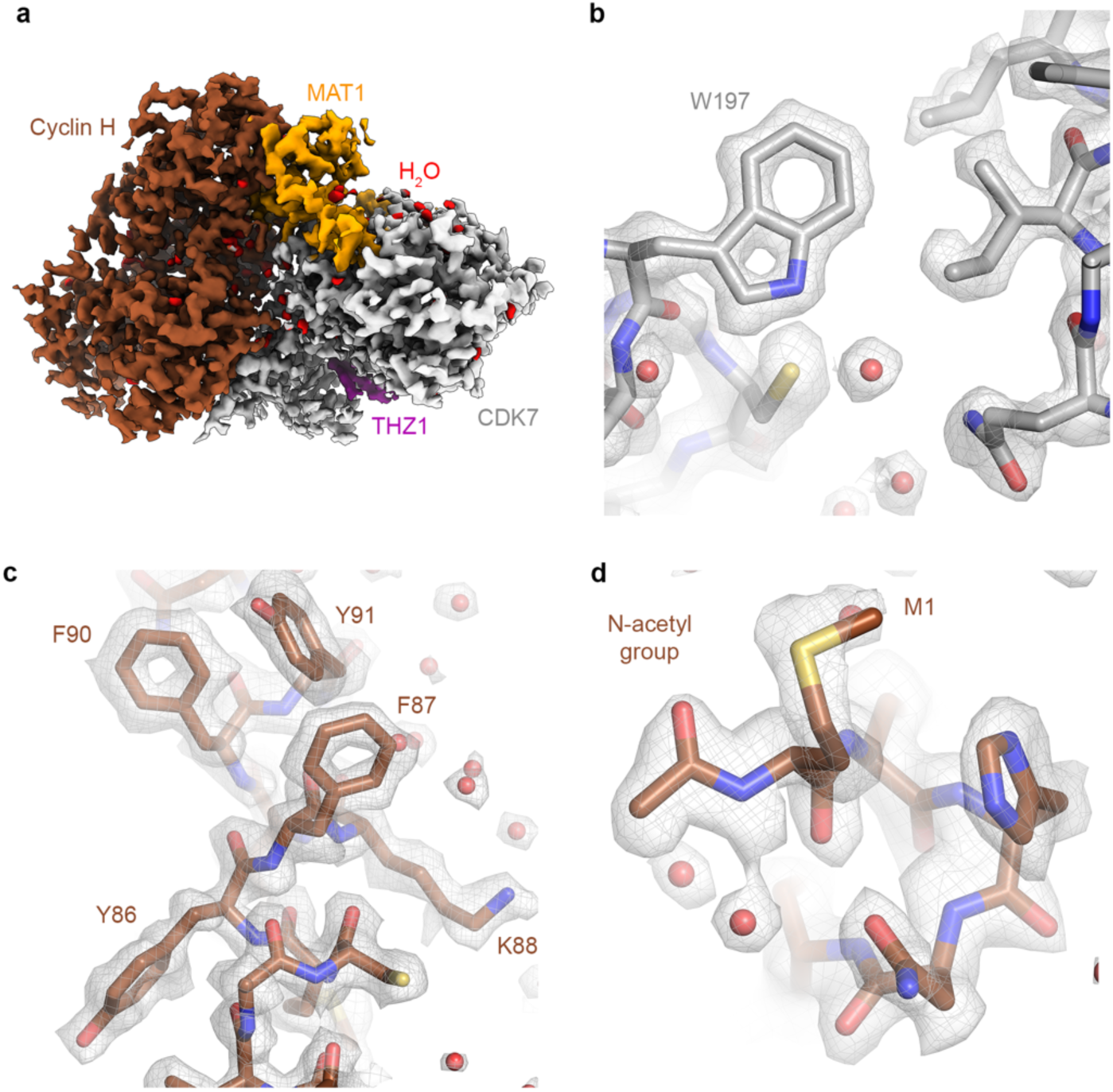
High-resolution structures of the human CDK-activating kinase. (**a**) Cryo-EM map with distinctly coloured CAK subunits (cyclin H brown, MAT1 orange, CDK7 grey, water molecules red, THZ1 purple). (**b**, **c**) Examples of high-quality density at 1.9 Å resolution (CAK- THZ1 map). (**d**) Direct identification of post-translational N-terminal acetyl modification on cyclin H in the cryo-EM map (CAK-ATPγS map).

The structure of CAK bound to the nucleotide analogue ATPγS at 1.9 Å resolution allows the accurate placement of the bound nucleotide and a water-coordinated Mg^2+^ ion in the active site (Extended Data Fig. 1f). Compared to the X-ray crystal structure of the homologous complex from *Chaetomium thermophilum*, the orientation of the adenosine base in the bound nucleotides is different (Extended Data Fig. 1g), highlighting the importance of using the human complex to support drug discovery. The structure of apo-CAK shows great similarity to the nucleotide- and inhibitor-bound structures, even for most of the active site pocket, except for the N-terminal β-sheet domain, which is substantially more flexible in the apo state than in all structures that have an occupied active site (Extended Data Fig. 1i-k). Flexibility limits the overall resolution of this reconstruction to 2.3 Å and is greatest at the outermost two β-strands, often referred to as the G-rich loop, which are barely visible in the density.

The structure of THZ1-bound CAK at 1.9 Å resolution (Fig. 1a, Extended Data Fig. 1d) agrees with our previous interpretation at lower resolution ^13^. Notably, even in our high-resolution map, the extended arm that carries the cysteine-reactive acrylamide warhead is resolved far

less well than the indole- and chloropyrimidine groups of the inhibitor that are embedded in the CDK7 active site pocket (Extended Data Fig. 1l). This is probably linked to flexibility of this part of the inhibitor, and it is unlikely that the resolution of this area could be improved with more data or by currently available image processing tools. We quantified these differences in local atom resolvability using Q-scores (Extended Data Fig. 1l) ^21^ because the local map quality in the presence of inhibitor heterogeneity does not necessarily closely correlate with the overall resolution of the reconstruction.

In contrast to THZ1, which acts via a covalent mechanism, LDC4297 is a highly selective competitive inhibitor of CDK7 ^3^. The map quality of our structure of CAK in complex with LDC4297 at 2.1 Å resolution (Extended Data Fig. 1e) was improved by locally refining only the approximately 40 kDa CDK7 density (Extended Data Fig. 1m). The structure reveals that the substituted phenyl ring assumes a similar conformation to the equivalent group in ICEC0942, while the ether-linked piperidine group assumes a boat-like conformation (Extended Data Fig. 1n).

### Cryo-EM workflow design and validation

To enable high-resolution and high-throughput cryo-EM data collection suitable for structure determination of larger numbers of small-molecule-bound CAK complexes, we devised and implemented a three-stage cryo-EM workflow comprising rapid initial grid and sample screening, intermediate-resolution structure determination, and finally high-resolution structure determination of the best grids (Fig. 2a). We applied this workflow to 12 compounds (Supplementary Table 1), but the approach can be scaled to even larger numbers of small-molecule ligands in the future.

**Figure 2.**
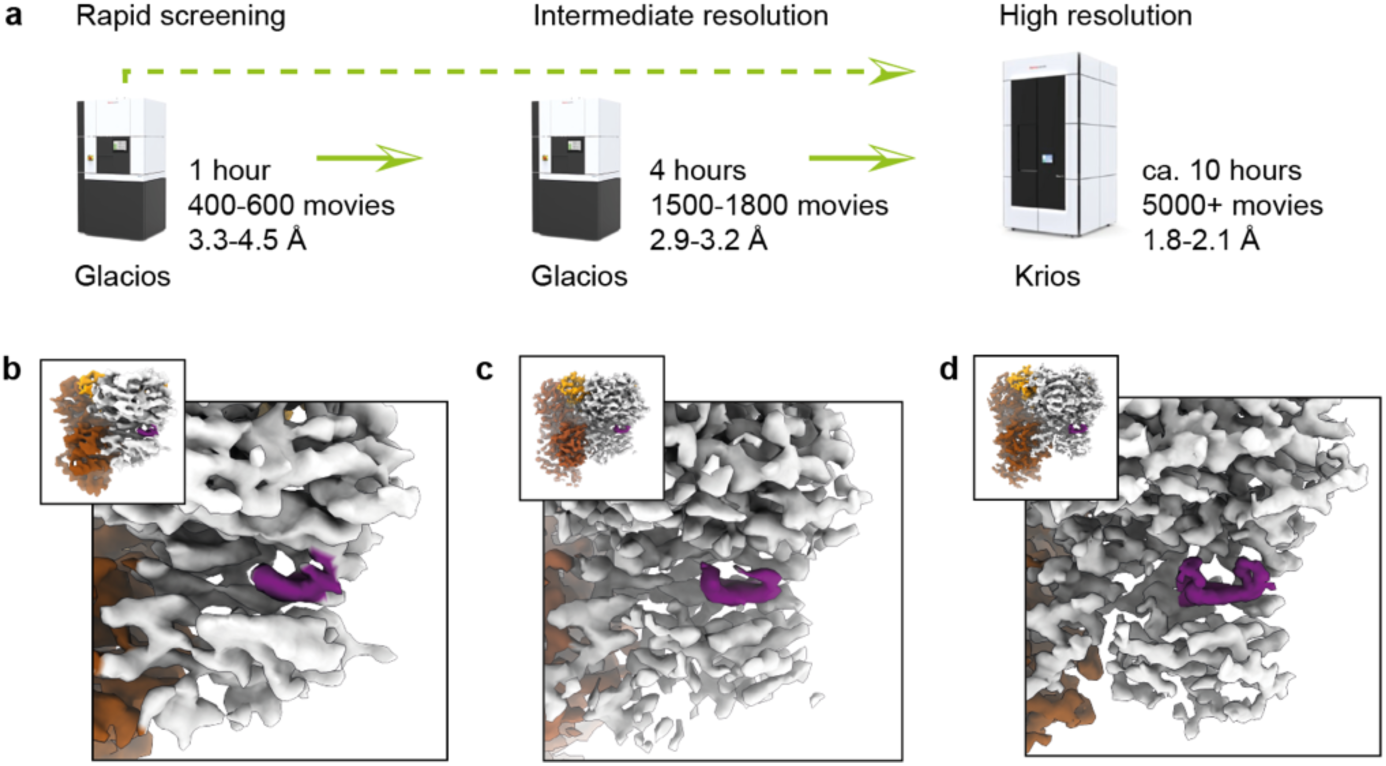
Cryo-EM workflow for inhibitor series structure determination. (**a**) Schematic of tripartite cryo-EM workflow. The possibility of using a two-stage workflow is indicated by a dashed green arrow. (**b**) Example result from 1-hour Glacios 2 screening. (**c**) Example result from 4-hour screening. (**d**) High-resolution cryo-EM map from Titan Krios G4 collection.

To aid in the design of this workflow, we first benchmarked the performance of a 300 kV Titan Krios G4 cryo-TEM equipped with a Falcon 4i detector, a Selectris X energy filter, and a cold-FEG against a 200 kV Glacios 2 cryo-TEM equipped with X-FEG, Falcon 4i detector, and Selectris X energy filter. We found that for the CDK-activating kinase, the 300 kV-cryo-TEM system provided an approximately 0.3 Å resolution advantage (Extended Data Fig. 2a-e), with resolutions of 2.0 Å and 2.3 Å achieved for data collected from the same grid, notably in a shorter time and from approximately 50% fewer electron micrograph movies for the 300 kV system compared to the 200 kV system (see Supplementary Note 1). We also investigated the contribution of the energy filter to high-resolution structure determination of our 85 kDa target complex. We found a 0.3 Å resolution loss in the absence of energy filtration under the conditions tested (Extended Data Fig. 2a, f, g, Supplementary Note 2). Given these results, we decided to use the 200 kV Glacios 2 setup for rapid sample screening and intermediate-resolution structure determination, and to leverage the 300 kV Titan Krios G4 setup for collection of high-quality datasets on selected specimens.

The goal of the initial rapid sample screening phase was the identification of specimens exhibiting (i) good ice thickness, (ii) suitable particle density, (iii) suitable particle orientation distribution, (iv) presence of inhibitor, and (v) promising resolution from on-the-fly processing of a dataset of a defined size during collection. We were able to satisfy these requirements using only 1 hour of data collection time per dataset (Fig. 2b), thereby determining 26 structures confirming the presence of 12 different bound inhibitors from 13 samples (Supplementary Table 1), most of them at approximately 3.5-4.0 Å resolution (Extended Data Figs 2h, 3, 4, Supplementary Table 1). All data processing at this stage was performed using cryoSPARC live ^22^, enabling rapid visualisation of the results during data collection.

**Figure 3.**
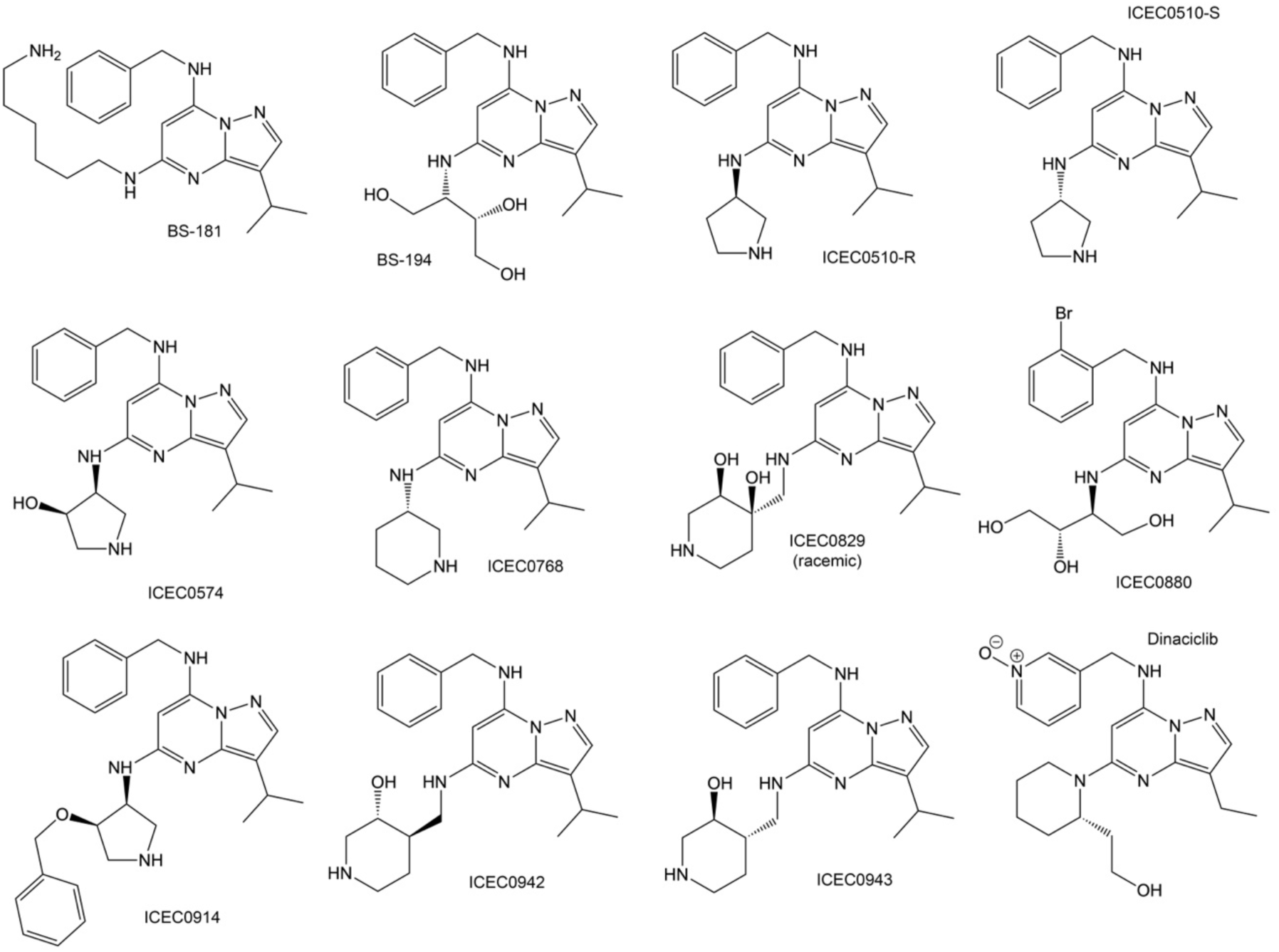
Chemical structures of pyrazolopyrimidine-type inhibitors analysed by high- resolution cryo-EM. See Extended Data Table 1 for enzyme inhibition properties of these compounds.

**Figure 4.**
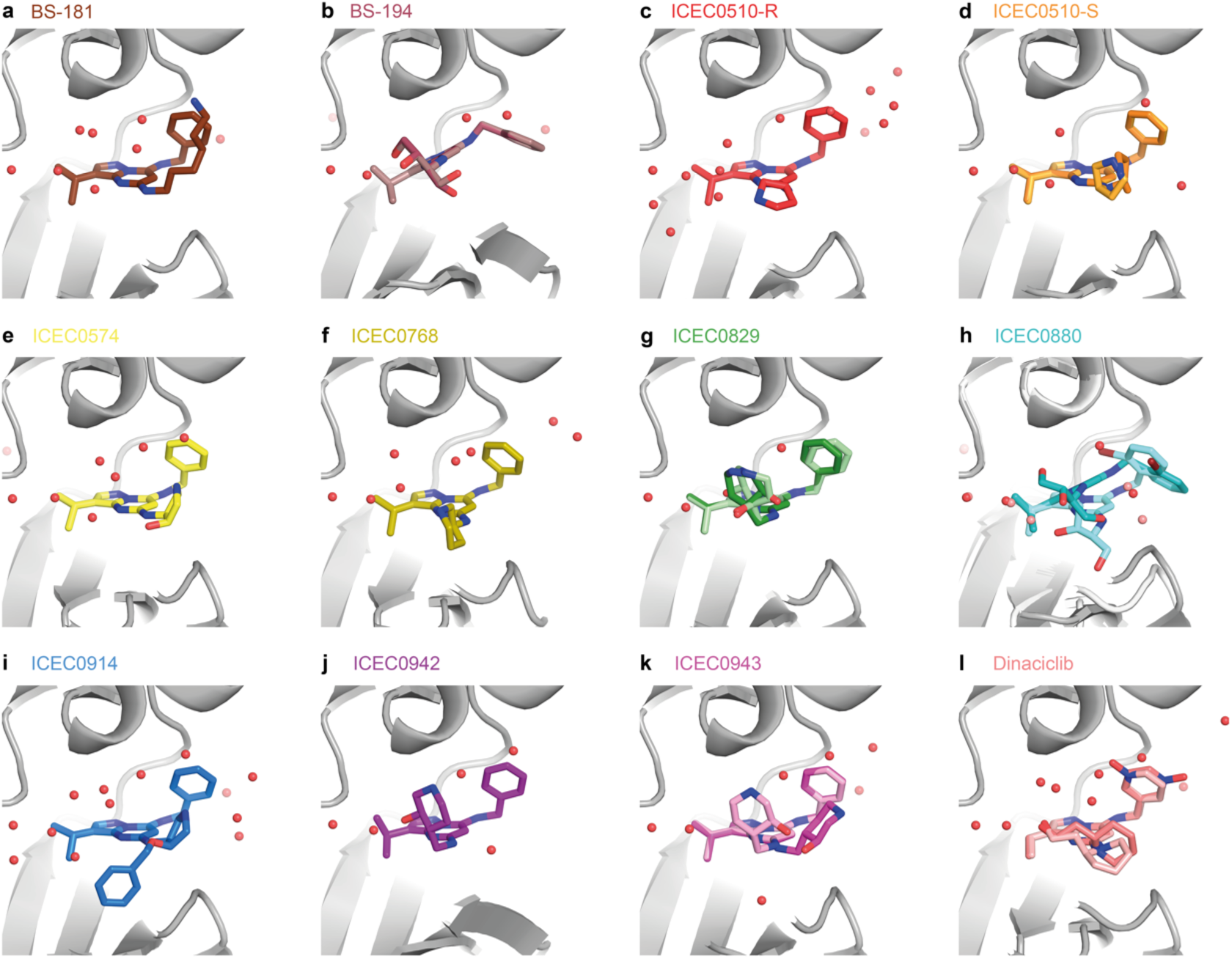
Structures of pyrazolopyrimidine-type inhibitors bound to CDK7. (**a**) BS-181. (**b**) BS-194, a compound selective for CDK2 (two conformers with minor differences). (**c**) ICEC510-R. (**d**) ICEC510-S (two conformers). (**e**) ICEC0574 (two conformers). (**f**) ICEC0768. (**g**) ICEC0829 (two enantiomers). (**h**) ICEC0880 (two different positions correlated with conformational changes in CDK7). (**i**) ICEC0914. (**j**) The clinical inhibitor ICEC0942. (**k**) ICEC0943, an enantiomer of ICEC0942 (two conformers). (**l**) Dinaciclib (two conformers).

We subsequently performed intermediate-resolution structure determination using the 200 kV Glacios 2 setup on selected grids, aiming to produce cryo-EM maps suitable for preliminary interpretation and approximate inhibitor docking. To this end, we extended the data collection time to 4 hours and achieved approximately 3 Å resolution for most datasets (Fig. 2c, Extended Data Fig. 2h, Supplementary Table 2). We applied a combination of cryoSPARC and RELION for processing of these datasets (see Methods for details). The complete processing of more than a dozen datasets was feasible in approximately one day after the end of data collection on a single 4-GPU server. Parallelisation of this step across multiple servers to reduce this time further will be straightforward if required. These acquisition sessions yielded 12 structures at around 3 Å resolution (Extended Data Fig. 5), in which inhibitors can be oriented and refined into the density and major conformational differences in large substituents are visible. Models derived from these reconstructions may also be used as starting points for molecular dynamics simulations or other computational methods, an option that we did not further pursue at this stage because we opted to collect high-resolution data for these complexes.

Aiming to resolve CAK-bound inhibitors at high resolution to provide highly accurate molecular models and identify water molecules that may contribute to inhibitor binding and specificity, we used the 300 kV Titan Krios G4 setup for high-resolution data collection (Fig. 2a, d). Data collections lasted for approximately 10 hours and yielded roughly 5,000 micrographs for each sample. These data were initially processed using cryoSPARC live and cryoSPARC ^22^ and final classifications, polishing, CTF refinement and 3D refinement were performed in RELION ^23^ . In contrast to the rapid screening and intermediate-resolution structure determination phases, the data processing effort for high-resolution reconstruction was substantial and extended substantially beyond the end of the data collection.

The large volume of CAK data acquired using this workflow (approximately 5,000 micrographs per dataset for 12 inhibitors, see below) also allowed us to explore the resolution limits for our specimen and instrument by combining 6 fully processed datasets that had previously reached 1.8-1.9 Å resolution individually (see below). From 2,060,503 particles, we obtained a CAK reconstruction with averaged inhibitor density at a resolution of 1.7 Å after CTF refinement and an additional round of particle polishing (Extended Data Fig. 6a-c). Given the large volume of data that entered these computations, further progress towards atomic-resolution structure determination of the CAK is likely to depend on new specimen preparation methods, such as the introduction of nanobody derivatives to improve alignability of the particle images ^24^, or on new instrumentation (Extended Data Fig. 6d, e).

### Structure determination of CAK-bound pyrazolopyrimidine-type inhibitors

ICEC0942 is an orally bioavailable high-affinity inhibitor of CDK7 that has entered clinical trials for cancer treatment ^4^. ICEC0942 was developed as a part of a series of pyrazolo[1,5- a]pyrimidine compounds. *In vitro* kinase inhibition data show that inhibitor selectivity spans almost 4 orders of magnitude, ranging from approximately 100x selectivity for CDK7 over CDK2 (ICEC0829) to 100x selectivity for CDK2 over CDK7 (BS-194) (Extended Data Table 1) ^4, 14, 25–28^, with low selectivity conferring a risk of inducing undesirable off-target effects in patients. Given the importance of understanding selectivity, and considering our previous observation of conformational differences of ICEC0942 bound to CDK7 and CDK2 ^12^, which suggests that structural analysis may aid in rationalising inhibitor selectivity and provide insights that may be applied to next-generation inhibitors, we undertook the high-resolution structural analysis of 12 pyrazolopyrimidine-type CDK inhibitors (Fig. 3) to explore the molecular mechanisms of CDK7 selectivity.

Our high-throughput screening and collection workflow enabled us to visualise 12 CAK- inhibitor complexes at 1.8-2.2 Å resolution (Fig. 4a-l, Extended Data Figs 6f-r, 7, Supplementary Table 3). The cryo-EM maps typically show well-resolved density for the pyrazolopyrimidine core and the benzylamine-derived substituents of the bound inhibitors, and less-well resolved density for the variable substituents at the C5 position of the pyrazolopyrimidine core, indicating structural heterogeneity in the bound compounds (e.g. Fig. 4g, k, l, Extended Data Fig. 7).

The high-resolution analysis of the CAK-ICEC0942 complex (Fig. 5a) extends and further defines our previous results, in which the orientation of the hydroxypiperidine substituent was ambiguous ^12^. Our high-resolution density supports an orientation of the six-membered piperidine ring in which the hydroxy group points into the active site cavity, where it hydrogen bonds with a water molecule that is in turn hydrogen bonded to CDK7 K41. Two additional water molecules are within hydrogen bonding distance of the piperidine nitrogen and the exocyclic amino group connecting the C5 substituent to the pyrazolopyrimidine core, respectively, and mediate interactions between the inhibitor and residues lining the CDK7 active site pocket (Fig. 5a). The hydroxy group on the ICEC0942 hydroxypiperidine substituent appears to be an important modulator of inhibitor binding, likely due to its ability to participate in the hydrogen bonding networks outlined above. The ICEC0942 stereoisomer ICEC0943 (Fig. 3) exhibits poorly defined density for the entire hydroxypiperidine ring, suggestive of two different orientations (Fig. 4k), explaining why it binds CDK7 with much reduced affinity, as reported previously ^14^.

**Figure 5.**
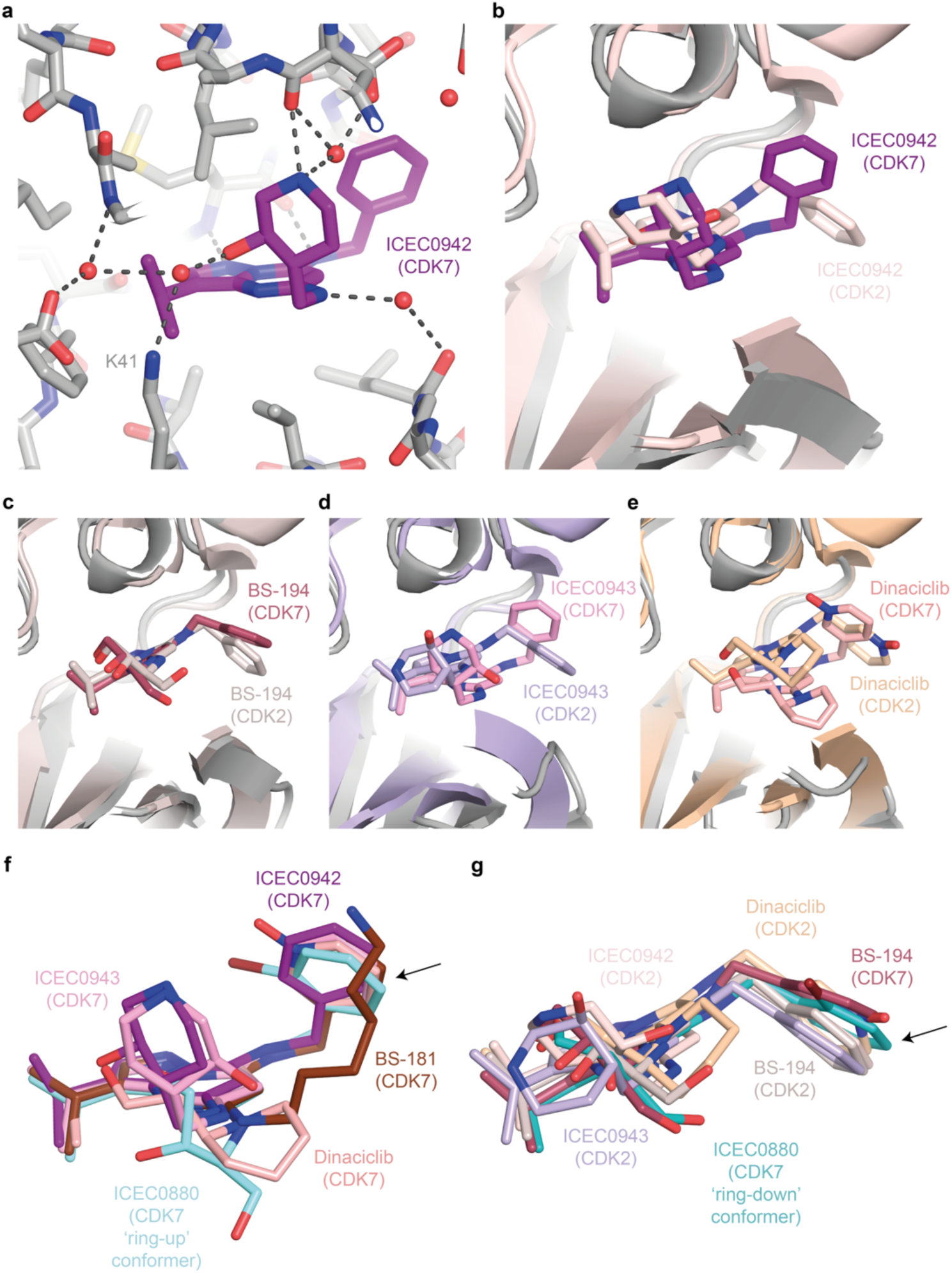
Analysis of high-resolution inhibitor structures. (**a**) Water molecules and hydrogen bonding network connecting ICEC0942 to surrounding residues in CDK7. (**b**) Comparison of ICEC0942 bound to CDK7 (this work) and CDK2 (PDB ID 5JQ5) ^14^. The conformation of the benzylamine group and the placement of the inhibitor core differ between the two structures. (**c**) Comparison of BS-194 bound to CDK7 (this work) and CDK2 (PDB ID 3NS9) ^26^. (**d**) Comparison of ICEC0943 bound to CDK7 (this work) and CDK2 (PDB ID 5JQ8) ^14^. (**e**) Comparison of dinaciclib bound to CDK7 (this work) and CDK2 (PDB ID 4KD1) ^29^. For clarity, only one conformer for the CDK7-bound compounds is shown in panels (c-e). (**f**) Superposition of ICEC0942, BS-181, ICEC0943, dinaciclib, and the ring-up ICEC0880 conformer, all bound to CDK7. (**g**) Superposition of ICEC0942, BS-194, ICEC0943, and dinaciclib, all bound to CDK2, with BS-194 and the ring-down conformer of ICEC0880 bound to CDK7. The flipping ring system is indicated with an arrow.

### Insights into CDK7 inhibitor selectivity

Our structural analysis yielded additional insight into inhibitor selectivity. BS-194 is the only inhibitor for which we observed a homogeneous “ring down” conformation of the benzylamine substituent at the C7 position of the pyrazolopyrimidine core (Fig. 4b). We observed previously that ICEC0942 assumes a “ring up” conformation bound to CDK7 and a “ring-down” conformation bound to CDK2 (Fig. 5b), suggesting a link to target selectivity ^12^. Comparisons of our new data with existing structures show that dinaciclib ^29^ and ICEC0943 ^14^ show a similar conformational switch between the CDK7 and CDK2-bound complexes, while BS-194 assumes a homogeneous ring-down conformation in complex with both kinases (Fig. 5c-e). Interestingly, BS-194 is strongly selective for CDK2 over CDK7, while the related derivative BS-181, which is selective for CDK7 over CDK2 (Extended Data Table 1) ^25, 26^, exhibits a “ring up” conformation bound to CDK7 (Fig. 4a, 5f). These observations strongly support our previous hypothesis that this conformational change is due to properties of the respective CDK active sites, and that the ability of an inhibitor to accommodate this change may be connected to target preference. The conformational change in the benzylamine ring is coupled to a shift of the pyrazolopyrimidine core of the inhibitors, which is observed for multiple compounds in CDK7 and CDK2 (Fig. 5b-g).

In contrast to the other inhibitors, which exhibit either clear “ring up” or “ring down” conformations, ICEC0880 exhibits both conformations when bound to CDK7 (Fig. 4h, 5f, g). Because these inhibitor binding modes coincide with movements of the N-terminal β-sheet domain of CDK7, it was possible to computationally separate the cryo-EM data into distinct classes and directly visualise and compare them (Fig. 4h, 5f, g). Given the small molecular weight of the inhibitor, which produces only a minute signal in the raw particle images, this would likely have been impossible in the absence of correlated motions in the protein.

It is worth noting that both BS-194 and its derivative ICEC0880 contain a 1,2,4-trihydroxybutyl substituent at the C5 position of their pyrazolopyrimidine core. It is possible that this substituent contributes to the observed effect by promoting a conformation of the pyrazolopyrimidine core that is conducive to the “ring down” conformation and reduced CDK7 selectivity (or increased CDK2 selectivity), which is counteracted by the bulky exocyclic bromine at the benzylamine substituent of the ICEC0880. This added steric constraint might destabilise binding in the “ring down” conformation, as exhibited by the shift of the entire pyrazolopyrimidine core in the two ICEC0880 structures, and it may contribute to the CDK7 selectivity of ICEC0880. Notably, LDC4297, which is highly specific for CDK7, carries an even more bulky substituent at its “ring up” benzylamine group (Extended Data Fig. 1b, k), and the recently reported CDK7-selective compound LGR6768 ^30^ exploits a bulky biphenyl substituent at this position.

## Discussion

We report a series of structures of the human CAK in free, nucleotide-bound, and small-molecule inhibitor bound states, 14 of them at 2 Å resolution or better. Notably, we obtained these results using a molecular complex that is the target of actively ongoing drug discovery programmes and clinical trials. Before our efforts, the structure of only one <100 kDa protein complex at better than 2 Å resolution had been deposited – that of streptavidin (EMD-31083) – and none for a medically relevant target or a protein complex lacking symmetry in this size range. We have now demonstrated that current cryo-EM technology is suitable for routine 2 Å structure determination of small protein complexes from less than one day of data collection on a high-end system, opening the door to high-throughput, high-resolution cryo-EM even for nominally difficult targets ^16^. Additionally, our screening approaches demonstrate the feasibility of the determination of more than a dozen 3.5-4.0 Å-structures a day, and several 3 Å structures a day, using a 200 kV microscope. Using a 300 kV instrument for screening harbours the potential for even higher sample throughput at the same resolution, or improved resolution at constant sample throughput. These results and the methodology outlined in this work will facilitate the application of cryo-EM to iterative structure-based drug design efforts, which are of critical importance to address global health challenges.

Our data show that the hydroxypiperidine substituent of the clinical inhibitor ICEC0942 engages in favourable interactions in the CDK7 active site, which results in the high affinity and selectivity of the compound. Our data additionally support a mechanistic hypothesis for a secondary mechanism of CDK7 selectivity in pyrazolopyrimidine-based CDK inhibitors, specifically that small structural differences in the nucleotide-binding pockets in CDK7 and CDK2 induce a preference for different conformations of the benzylamine substituent that forms part of the scaffold of the inhibitors we analysed, and that chemical modifications that favour one conformation over the other may be used to improve kinase selectivity of these compounds. These findings will pave the way for future structure guided CDK7 inhibitor design. Furthermore, they provide a proof of principle for the benefit of the application of cryo-EM in structure-based CDK7 inhibitor design.

## Methods

### Synthesis of compounds

Pyrazolopyrimidine compounds were synthesised according to published strategies ^4, 25–27^. All characterisation data were consistent with prior studies. The identities and purities of all compounds were confirmed by UV-vis spectrometry and mass spectrometry, as described below. All compounds were shown to be >96 % pure prior to biological studies. ATPγS, THZ1 and LDC4297 were purchased commercially from Sigma Aldrich, EMD Millipore, and MedChemExpress (distributed by Insight Biotechnology, Wembley, UK), respectively.

### Verification of inhibitor integrity and purity

LC/MS grade solvents, formic acid, or alternative eluent modifiers were purchased from VWR (Poole, UK) and Fisher (Loughborough, UK) unless otherwise stated. 10 μL of 1 mM DMSO solution of each compound was plated in a 384 Greiner (781280) well plate. 0.1 μL standard injections (with needle wash) of the sample were made onto a Phenomenex Kinetex C18 30 x 2.1 mm, 2.6u, 100A column (Phenomenex, Torrance, CA, USA). Chromatographic separation was carried out at 40°C using an Agilent 1260 Infinity II series UPLC (Agilent, Santa Clara, USA) over a 4 minute gradient elution (KNOWNS_AM190319.m) from 90:10 to 10:90 water:methanol (both modified with 0.1% formic acid) at a flow rate of 0.4 mL/min. UV-vis spectra were acquired at 254 nm on a 1260 Series diode array detector (Agilent, Santa Clara, USA).

The post column eluent flow from the diode array detector was split, with 90% sent to waste. The remainder was infused into a 6530 Series QtoF mass spectrometer fitted with an Agilent Jet Stream ESI source (Agilent, Santa Clara, USA). LC eluent and nebulising gas was introduced into the grounded nebuliser with spray direction orthogonal to the capillary axis. A nozzle voltage of 0 V was applied to the charging electrode to generate a charged aerosol. The aerosol was dried by heated drying gas (10 L/min of nitrogen at 350°C, 35 psi), producing ions by ESI. Ions entered the transfer capillary along which a potential difference of 4 kV was applied. The fragmentor voltage was set at 175 V and skimmer at 65 V. Signal was optimised by SWARM autotune. Profile mass spectrometry data was acquired in positive ionisation mode over a scan range of m/z 190 – 650 (scan rate 4.0) with reference mass correction at m/z 622.02896 hexakis(2,2-difluroethoxy)phosphazene.

Raw data were processed using Agilent MassHunter Qualitative Analysis B.07.00 (AutoQC.m). The “Find Compounds by Formula” algorithm was used to identify compounds and calculate the purity. The compound purity was calculated using the highest value of %UV (at 254 nm) or %TIC (total ion count from the mass spectrometer).

### Enzyme inhibition assays

In-vitro kinase assays of inhibitors were performed in ProQinase format as described ^4, 14^.

### CAK expression, purification, and cryo-EM specimen preparation

CAK was expressed in insect cells and purified by nickel-affinity, strep-tactin, and gel filtration chromatography as described ^13^. Most datasets used the CAK-MAT1Δ219 variant, in which the N-terminal 219 residues of MAT1 are replaced by an MBP-tag, which is then cleaved by incubation with TEV protease, as described previously ^13^. The LDC4297 dataset additionally used the CAK-peptide construct, also described previously ^13^.

Cryo-EM specimens were prepared on UltrAuFoil R1.2/1.3 holey gold grids (Quantifoil Microtools). For complex formation, CAK complex at approximately 2 mg/mL stock concentration was diluted 5-6x in sample buffer (20 mM Hepes-KOH pH 7.9, 200 mM KCl, 2 mM MgCl^2^, 5 mM β-mercaptoethanol) and incubated with 50 µM inhibitor (dissolved at 50 mM concentration in 100% DMSO) or 2 mM ATPγS (Merck Life Science UK) at room temperature for 5 min. Covalent CAK-THZ1 complex was prepared as described ^13^. 4 µL of CAK complex were applied to a plasma cleaned grid (Tergeo plasma cleaner, PIE scientific) mounted in a Vitrobot Mark IV (Thermo Fisher Scientific) operated at 5°C and 100% humidity, blotted for 1, 1.5, 1.5, or 2 sec (4 grids made for each complex) and plunged into liquid ethane at liquid N^2^ temperature. After vitrification, grids were clipped into autogrid cartridges (Thermo Fisher Scientific) for use with Glacios and Titan Krios autoloader systems.

### Initial screening

Initially, grids were coarsely screened on a Glacios cryo-transmission electron microscope (cryo-TEM) operated at 200 kV acceleration voltage and equipped with a Falcon 4 (later Falcon 4i) direct electron detector. Data were acquired in EER format at a pixel size of 0.5675 Å/pixel using a flux of 4-6 electrons . pixel^-1^ . second^-1^ using aberration-free image shift (AFIS) to accelerate the data collection. Approximately 300-600 micrographs were processed in real time using cryoSPARC live ^22^ using 2x binning and with exposures fractionated into 60 frames for motion correction, which provided results suitable to judge overall sample quality and particle orientation distribution. This approach was later supplanted by the rapid-screening approach on a Glacios 2 electron microscope equipped with a Falcon 4i detector, a Selectris X energy filter, and fringe-free imaging capability enabling collection of 2-3 movies per hole and increased data collection rates. The latter data are shown in the Results section.

### Multigrid-screening and data processing

We used acquisition settings that were similar to our previously reported conditions for structure determination of the CAK ^12, 13^, adjusted for the changes in hardware compared to prior experiments, resulting in 0.57 Å pixel size, 70 electrons . Å^-2^ total exposure at 7-8 electrons . pixel^-1^ . sec^-1^ and EER format fractionated into 50 frames for screening runs and 70 frames for high-resolution runs (see below).

Data were collected using EPU-Multigrid on a 200kV Glacios 2 microscope equipped with a Selectris X energy filter coupled with a Falcon 4i detector (Thermo Fisher Scientific). Data were collected using fringe free imaging (FFI) and AFIS at a throughput of roughly 500 images/hr. 2 grids each of 5 different samples and a dry Quantifoil Cu200 R2/2 or S106 cross-grating (11 grids total) were loaded into the autoloader for each EPU-Multigrid run. Within EPU, a new session queue was created, followed by sequentially loading grids to the stage for session set- up and grid square selection to give approximately 500 (for 1hr-screening runs) or 2000 (for 4-hour data collection session) images after automatic ice-filter selection. Before starting data collection, two-fold astigmatism was corrected, and beam tilt was adjusted to the coma-free axis using using the dry Quantifoil Cu200 R2/2 or S106 cross-grating. The Selectris X filter slit was then centred on the zero-loss peak with a slit width of 10eV, and the filter tuned for isochromaticity, magnification and chromatic aberrations using Sherpa software (Thermo Fisher Scientific) over vacuum. The EPU Multigrid queue was then started, and each grid was automatically loaded sequentially onto the stage, whereupon grid squares previously selected were automatically brought to eucentric height and holes for data collection were automatically selected with ice filter settings. Images were collected at a nominal magnification of 205,000x with a resulting pixel size of 0.57 Å. Flux measured on the detector over vacuum at spot size 3 at parallel illumination conditions was 7.94 electrons . pixel^-1 .^ sec^- 1^ and each exposure was 2.85 sec long to accumulate a total dose of 70 electrons . Å^2^ per exposure. Images were collected at defocus values -0.8µm, -1.0µm and -1.2µm. AFIS together with FFI and a 20 µm C2 aperture was employed to acquire 2 exposures per 1.2 µm ice hole and to accelerate data collection to approximately 500 per hour.

Rapid-screening and intermediate-resolution data were processed in cryoSPARC live. Movies were binned 2x during motion correction. Particles were picked using blob picker (elliptical blob, 80-110 Å diameter) and extracted in 160 x 160-pixel boxes. On-the-fly 2D classification and 3D refinement were performed using default parameters, with a previous reconstruction of CAK serving as an initial reference. Using a workstation with 4 RTX3090Ti GPUs (Nvidia Corporation), the software was able match data collection rates.

The rapid screening workflow used cryoSPARC live processing exclusively. To improve interpretability of the maps in the intermediate resolution workflow, we added two steps after cryoSPARC live processing: (i) We performed an additional 2D classification of all live- picked particles (except obvious non-particles and artifacts) which led to recovery of more particles and clear classes for additional particle views. Subsequently (ii), we performed a 3D refinement, an alignment-free 3D classification, and a final refinement of the best class in RELION ^23^, providing the final reconstruction. The resolutions were typically almost unchanged (within 0.1 Å) compared to the cryoSPARC live output, but the RELION maps were cleaner. The cryoSPARC 2D classification took approximately 2 hours per dataset, and the downstream processing in RELION was completed in approximately 1.5 hours.

### High-resolution data collection

Data were collected on a Titan Krios G4 cryo-TEM (Thermo Fisher Scientific) equipped with a cold-FEG operating at 300 kV acceleration voltage, a Selectris X energy filter, and a Falcon 4i direct electron detector. Data were acquired at 0.57 Å pixel size, 70 electrons . Å^-2^ total exposure at 7-8 electrons . pixel^-1^ . sec^-1^ and in EER format, later fractionated into 70 frames during data processing. AFIS and FFI were employed to acquire 2-3 exposures per 1.2 µm ice hole and to accelerate data collection to approximately 600 movies/hour by collecting data in holes within 12 µm of image-beam shift radius from the centred position. Approximately 5000 images were collected per grid for most grids (10,000 for THZ1).

### Pre-processing of high-resolution datasets in cryoSPARC

*General pre-processing strategy*: Datasets were acquired and processed in cryoSPARC live ^22^ with the same settings as for the screening datasets (see above). In addition to blob picking during on-the-fly processing, two parallel template-based picking strategies were then carried out after the end of the data collection, using representative 2D classes from the corresponding live-processing session or 2D projections generated from a 3D reconstruction as templates (Extended Data Fig. 9). Template-based picks were separately extracted from the micrographs “accepted” during live processing, using an extraction box size of 160 x 160 pixels, and classified into 200 2D classes (100-140 iterations, circular mask diameter 104-110 Å, batch size per class 200). For most datasets, the best 2D classes from the live processing session (blob picking) and from the two post-acquisition template-based picking strategies were selected, combined and duplicate particles were removed. The combined particles were re-extracted to improve particle centring and any duplicate particles arising from the re-centring during re-extraction were removed again. This final particle set was exported and converted to a *.star file compatible with RELION using a conversion program available in the PYEM package ^31^ followed by adjustment of path names to adhere to RELION conventions. This strategy was applied to all datasets with occasional small variations, outlined in the paragraphs below.

*CAK-THZ1*: For each of the two CAK-THZ1 datasets, which were collected early and used for trialling of processing strategies, the particles selected from the three picking strategies (1x blob picking, 2x template picking) were separately re-extracted before combination into one dataset, duplicate particle removal, and export to RELION format (“combined particles”). Additionally, the particles selected from blob picking and on-the-fly 2D classification in cryoSPARC live were exported as well (“blob particles”). The “combined particles” and the “blob particles” were separately imported into RELION, giving a total of four imported particle sets (two from each CAK-THZ1 grid).

*Apo-CAK*: For the apo-CAK complex, particles were additionally classified into 50 2D classes after the first duplicates removal step to remove any classes with concentric ring patterns arising from sensor imperfections from the selected particle set.

*CAK-ICEC0510-S*: the final duplicate particle removal step occurred within RELION after a 3D refinement.

### High-resolution data processing in RELION

*General refinement strategy*: All datasets were further processed using RELION 4.0 (Extended Data Fig. 8). To ensure full RELION functionality, including Bayesian polishing ^32^, movies were motion corrected using the RELION CPU-based implementation of the motion correction algorithm ^33^, using a binning factor of 2.0 (giving a pixel size of 1.14 Å/pixel). Exported particles from cryoSPARC were re-extracted from the motion corrected micrographs using an extraction box size of 160 x 160 pixels. Extracted particles were subjected to masked 3D refinement using a previous CAK reconstruction as an initial reference ^13^, with an initial angular sampling of 7.5 degrees. For ligand-bound datasets, the refined particles were classified into four 3D classes by alignment-free 3D classification (τ = 24, mask diameter 104 Å, resolution E-step limited to 4 Å). The best class(es) were selected, refined, and subjected to Bayesian polishing ^32^, after first training the polishing on 10,000 particles, with re- windowing and re-scaling to a box size of 384 pixels at 0.855 Å/pixel to account for CTF delocalisation. Polished particles were refined and subjected to CTF refinement of per-particle defocus followed by refinement of beam tilt, trefoil, and fourth-order aberrations or vice versa (depending on which strategy provided the best result for each dataset), followed by 3D refinement and a second round of polishing with re-windowing and re-scaling to a box size of 480 pixels at 0.7125 Å/pixel. Re-polished particles were re-refined and the resulting map was post-processed in RELION. Deviations from this general strategy are outlined in the following paragraphs.

*CAK-ATPγS*: After completion of the processing pipeline outlined above, a final 3D classification step using 2 classes (τ = 36) was inserted to identify the best-quality particle set used for the final refinement.

*CAK-LDC4297*: Due to sample heterogeneity, preferred orientation, and the presence of non-particle images, two additional 2D classification steps were inserted between the first 3D classification and the first Bayesian polishing step, selecting approximately 61,000 rare particles views to be added to the best 3D class, and eliminating non-particles from the resulting dataset. Subsequent polishing, refinements, and CTF refinements were performed according to the scheme outlined above. As a final step, a local refinement using a mask encompassing only CDK7 was performed to improve the appearance of the density of the inhibitor (Extended Data Fig. 1j, k).

*CAK-BS-181, CAK-ICEC0942 (grid VC13-3), CAK-ICEC0943*: The training step of Bayesian polishing failed, and particle polishing was instead run using the following *ad hoc* parameters, chosen based on the results of successful training runs from other samples, which used the same microscope and identical grid type: α_vel_ = 1.2, α_div_ = 4500, and α_acc_ = 1.5.

*Apo-CAK*: Training for Bayesian polishing failed, and polishing was run with *ad hoc* parameters (as described above). Due to conformational heterogeneity of the β-sheet domain in apo- CAK, particles were 3D classified while masking out the β-sheet domain after the initial motion correction, particle extraction, and initial refinement steps. This was done to focus the classification on the bulk of the particle density to select for particle quality, rather than letting the mobile β-sheet density dominate the classification. The selected particles were then refined using a full mask and polished only once (with re-windowing and re-scaling to a box size of 384 pixels at 0.855 Å/pixel) before a final refinement.

*CAK-THZ1*: Each of the two datasets (grids BG29-1, BG29-2) was motion corrected and each of the four exported particle sets from cryoSPARC (see above) were separately extracted from their corresponding micrographs. Extracted particles were separately refined and 3D classified, before selected particles were combined, duplicates were removed, and the resulting particle set refined before the initial polishing step. Training for Bayesian polishing failed, and polishing was run with *ad hoc* parameters (as described above). After the second polishing step, refined particles were additionally classified into 96 2D classes (τ = 2, mask diameter 110 Å, E-step resolution limited to 6 Å) to remove non-particle classes before the final refinement.

*CAK-ICEC0510-S, CAK-dinaciclib*: After the first 3D classification step, particles were selected from two different iterations (best class from iteration 15, two best classes from iteration 25), refined separately, joined, and subjected to duplicate removal. The resulting particle set was refined and polished (using *ad hoc* parameters because training failed, see above). Polishing and re-refinement steps were carried out according to the standard procedure above, but a 3D classification (3 classes) was inserted between CTF refinement and the second polishing run to improve the quality of the final particle set.

*CAK-ICEC0574*: CTF refinement consisted of aberrations followed by per-micrograph astigmatism.

*CAK-ICEC0768*: CTF refinement included only aberrations (beam tilt, trefoil, 4^th^ order aberrations) and an additional 3D classification (3 classes, τ = 24) was performed after the two polishing runs to identify the final particle set.

*CAK-ICEC0880, ICEC0829*: Additional 10-class 3D classifications (without resolution E-step limit) were carried out following the second round of polishing for ICEC0880 and ICEC0829. For ICEC0880, this revealed two distinct inhibitor conformations; classes corresponding to these two conformations were selected and refined. For ICEC0829, classes with the clearest inhibitor density were selected, combined, and refined.

*CAK-ICEC0942* (grid VC14-1): An additional 3D classification step (4 classes, τ = 24) after the first round of polishing and three rounds of CTF refinement (initially only refining beam tilt and third order aberrations, then defocus, then beam tilt and all aberrations) were performed.

### Model building and refinement

The CAK-THZ1 model was generated by rigid-body fitting of the previously published model ^13^ (PDB accession code 6XD3) into the post-processed map in UCSF ChimeraX ^34^. This was followed by several rounds of manual rebuilding in COOT ^35^, during which water molecules were manually modelled based on inspection of the density and hydrogen bond distances, and real-space refinement in PHENIX ^36^. The refined CAK-THZ1 model was used as a starting point for building the remaining models, for which ligands and restraint files were generated using PHENIX eLBOW ^37^. Ligands were manually modelled into the densities in COOT and models were adjusted and refined in the same way as for CAK-THZ1. For CAK-LDC4297 and CAK-ICEC0942, the structure was additionally refined using the PHENIX-OPLS4 pipeline ^38^ using the SCHRODINGER 2021-4 software package, followed by re-refinement in PHENIX while using reference restraints on the ligand, which resulted in improved overall MOLPROBITY score and model-to-map fit. For CAK-ICEC0880 (“ring-up” conformer), the structure was refined using reference restraints on CDK7 residue E68, resulting in an improved model-to-map fit of a nearby water molecule. Water molecules with Q-score < 0.70 were excluded from the models, except in the case of the water that is hydrogen bonded to N26 of the ligand in the CAK-ICEC0574 structure, which has a Q-score of 0.69 but whose presence is clearly supported by the density and the presence of equivalent water molecules in several other structures. Refinement statistics are provided in Supplementary Tables 4-10.

## Data availability

Atomic coordinates of high-resolution ligand-bound complexes have been deposited to the Protein Data Bank (PDB) using identifiers XXXX, XXXX, XXXX, XXXX, XXXX, XXXX, XXXX, XXXX, XXXX, XXXX, XXXX, XXXX, XXXX, XXXX, XXXX, XXXX, XXXX, XXXX, XXXX, XXXX, and XXXX. High-resolution cryo-EM maps have been deposited to the Electron Microscopy Data Bank (EMDB) using accession codes XXXX, XXXX, XXXX, XXXX, XXXX, XXXX, XXXX, XXXX, XXXX, XXXX, XXXX, XXXX, XXXX, XXXX, XXXX, XXXX, XXXX, XXXX, XXXX, XXXX, and XXXX. Electron micrograph movies for selected datasets have been deposited to the Electron Microscopy Public Image Archive (EMPIAR) with accession codes XXXX, XXXX, XXXX, XXXX, XXXX, XXXX, XXXX, and XXXX.

## Acknowledgments

We thank Fabienne Beuron for help with electron microscopy, Chris Richardson for help with high-performance computing, Gary Newton for providing the OPLS4 computing environment, and Claudio Alfieri for providing access to his GPU servers. We thank Meirion Richards, Amin Mirza and the ICR Structural Chemistry team for LC-MS verification of compound identity and integrity. B.J.G. was supported by a career development fellowship from the Medical Research Council of the UK (MR/V009354/1) and V.I.C. was funded by an ICR PhD studentship. S.A. received funding by Cancer Research UK grants C37/A9335, C37/ A12011, and C37/A18784. K.J. was supported by a PhD studentship from OneMoreCity and Imperial College London.

## Authors Contributions

B.J.G, A.K., and S.A. designed the cryo-EM-based project. K.J. characterised CAK inhibitors. B.J.G., V.I.C., and J.F. prepared the cryo-EM specimens and performed initial screening. A.F.K. and A.K. collected all cryo-EM data contained in the manuscript. V.I.C. and B.J.G. processed the cryo-EM data. V.I.C. interpreted the cryo-EM data under the guidance of B.J.G. B.J.G. and V.I.C drafted the first version of the manuscript, and all authors contributed to its final form.

## Competing interests

A.K. and A.F.K are employees of Thermo Fisher Scientific, the manufacturer of the electron microscopes used in this study. S.A. is a named inventor on patents concerning CDK7 inhibitors, which have been licensed to Carrick Therapeutics, and he owns shares in Carrick Therapeutics. A.K.B. is an employee and shareholder of Carrick Therapeutics.

## Extended Data Figure Legends

**Extended Data Figure 1.**
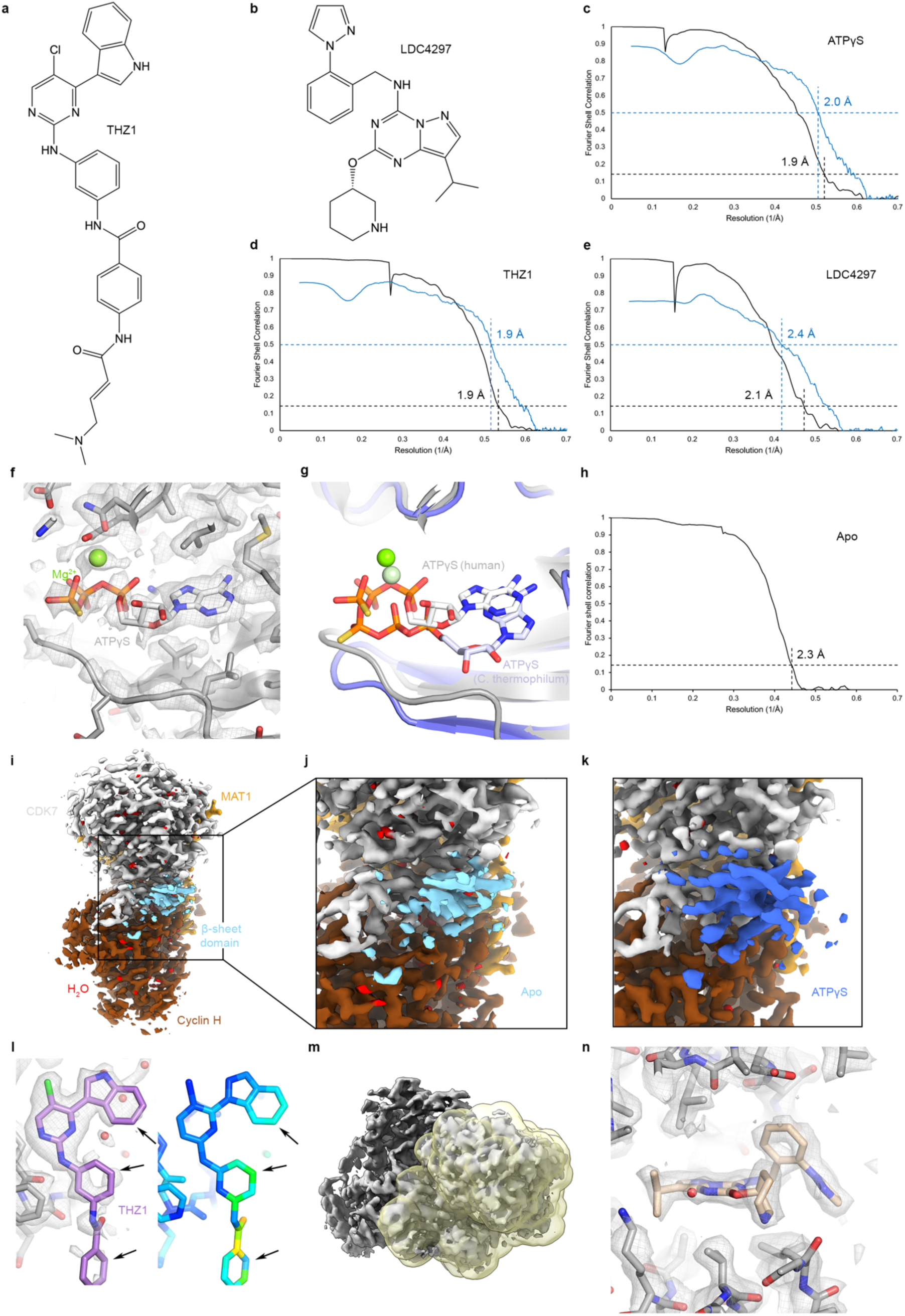
Structures of apo-CAK and CAK with bound nucleotide, THZ1, and LDC4297. (**a**) Chemical structure of THZ1. (**b**) Chemical structure of LDC4297. (**c**-**e**) Fourier shell correlation (FSC) curves for the CAK-ATPγS, CAK-THZ1, and CAK-LDC4297 cryo-EM reconstructions. Half-map FSC curves are shown in black, model vs. map FSC curves in blue. Resolutions are estimated according to the FSC = 0.143 criterion for half-maps, and the FSC = 0.5 criterion for model vs. map FSCs ^39^. (**f**) Structure of the ATPγS-bound active site of CAK shown with the cryo-EM map. (**g**) ATPγS molecule bound in the active site of human CDK7 (white) compared to adenosine nucleotide bound in the active site of fungal CDK7 (PDB ID 6Z4M; light blue). (**h**) FSC curve for the apo-CAK cryo-EM reconstruction. (**i**) Cryo-EM map of apo-CAK, with β-sheet domain shown in cyan. (**j**, **k**) Comparison of the density for the β-sheet domain in nucleotide-bound and apo-CAK (domain highlighted in cyan and blue, respectively; both maps low-pass filtered to 2.3 Å resolution). (**l**) THZ1 head group occupying the active site of human CDK7 (left). The cryo-EM map reflects the heterogeneity of less-tightly bound components of the inhibitor, which is quantified using Q-scores (right). (**m**) Mask used for masked refinement of a 40 kDa-fragment shown in yellow, CAK in grey). (**n**) Anti-viral compound LDC4297 bound in the active site of CDK7.

**Extended Data Figure 2.**
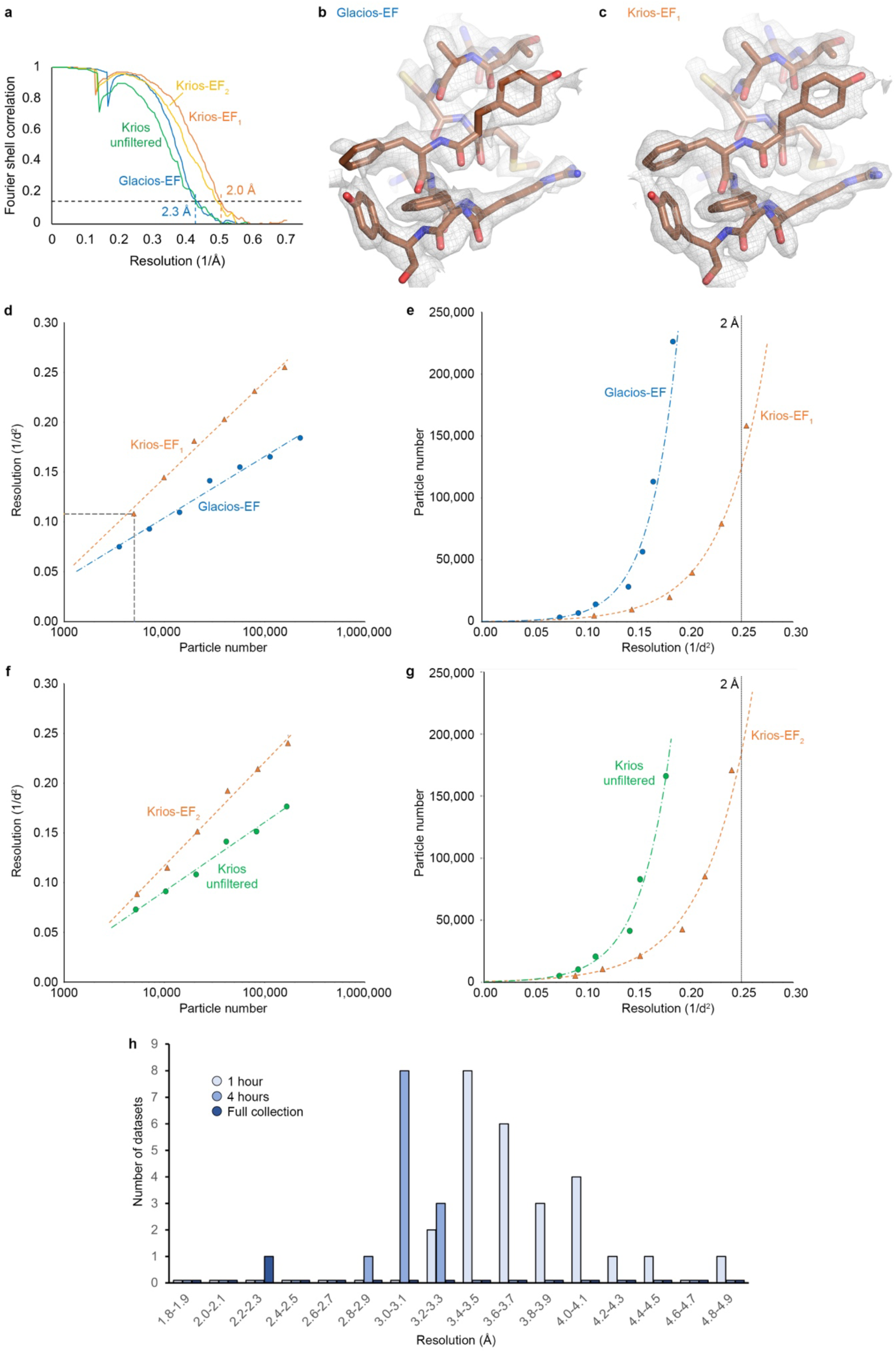
Glacios vs. Titan Krios resolution comparison and impact of energy filtration on the resolution obtained. (**a**) FSC curves for the Glacios and Krios cryo-EM reconstructions (EF = energy filtered). (**b**, **c**) Comparison between cryo-EM maps from Glacios- EF (b) and Krios-EF_1_ (c) data. (**d**) Henderson-Rosenthal plot comparing Krios-EF_1_ and Glacios- EF data. The inverse squared resolution of data subsets is plotted against the logarithm of the particle number, resulting in a linear relationship. (**e**) Particle number plotted against inverse squared resolution. It is apparent that the Glacios data may not approach the 2 Å line even for very large amounts of data. (**f**, **g**) As d and e, but comparing the reconstructions obtained from the energy-filtered Titan Krios setup (dataset Krios-EF_2_) against the results obtained after retracting the slit from the beam path (Krios unfiltered). (**h**) Overview of resolutions achieved in the Glacios data collections.

**Extended Data Figure 3.**
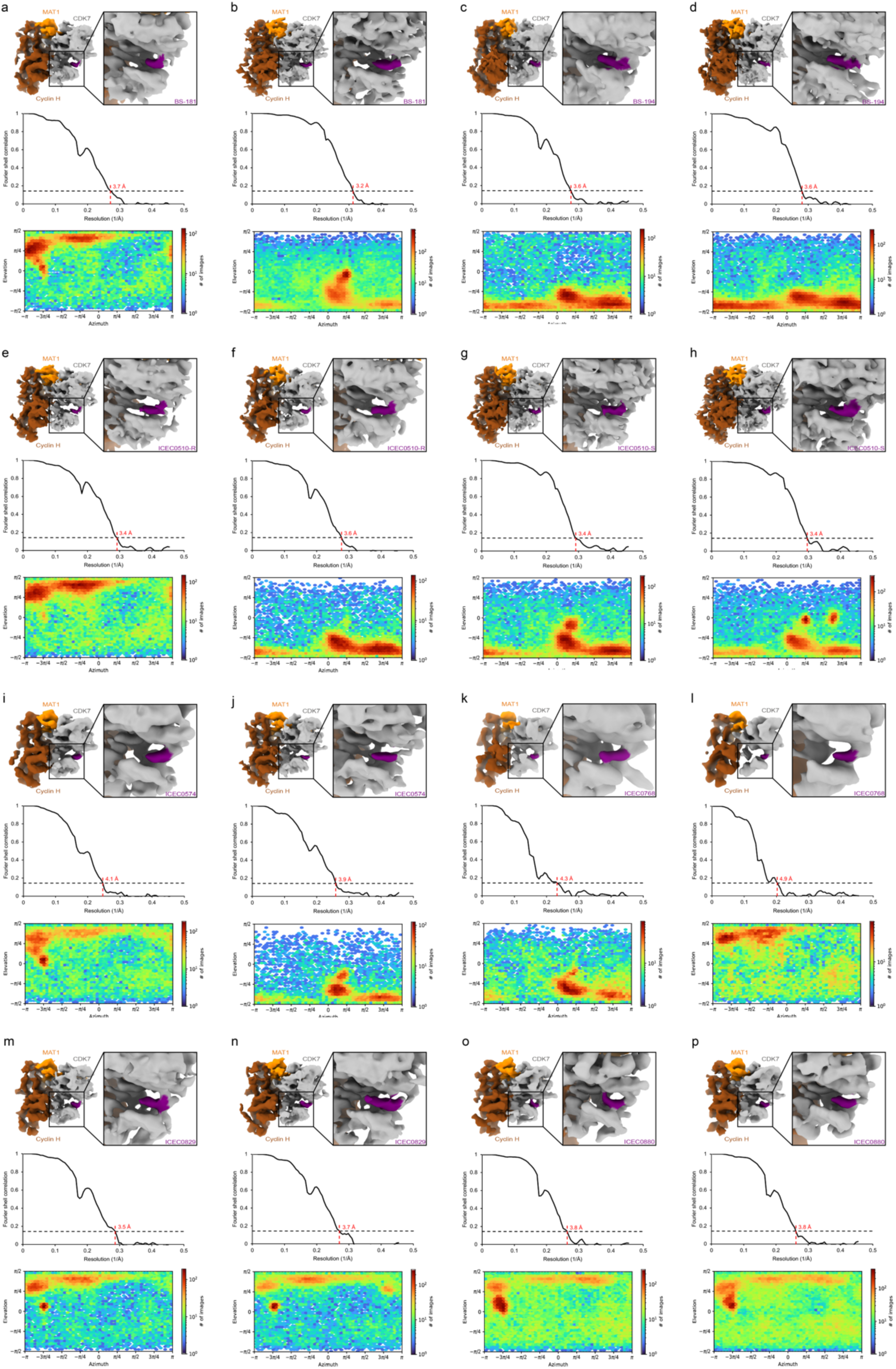
Live processing of 1-hour Glacios datasets, part 1. (**a**-**p**) Overview of 1-hour Glacios datasets. Top panels show a view of the 3D reconstruction (CDK7 grey, cyclin H brown, MAT1 orange) and a close-up view of the density for bound inhibitors (purple). Middle panels show the resolution according to the FSC = 0.143 threshold. Bottom panels show orientation distribution plots from cryoSPARC.

**Extended Data Figure 4.**
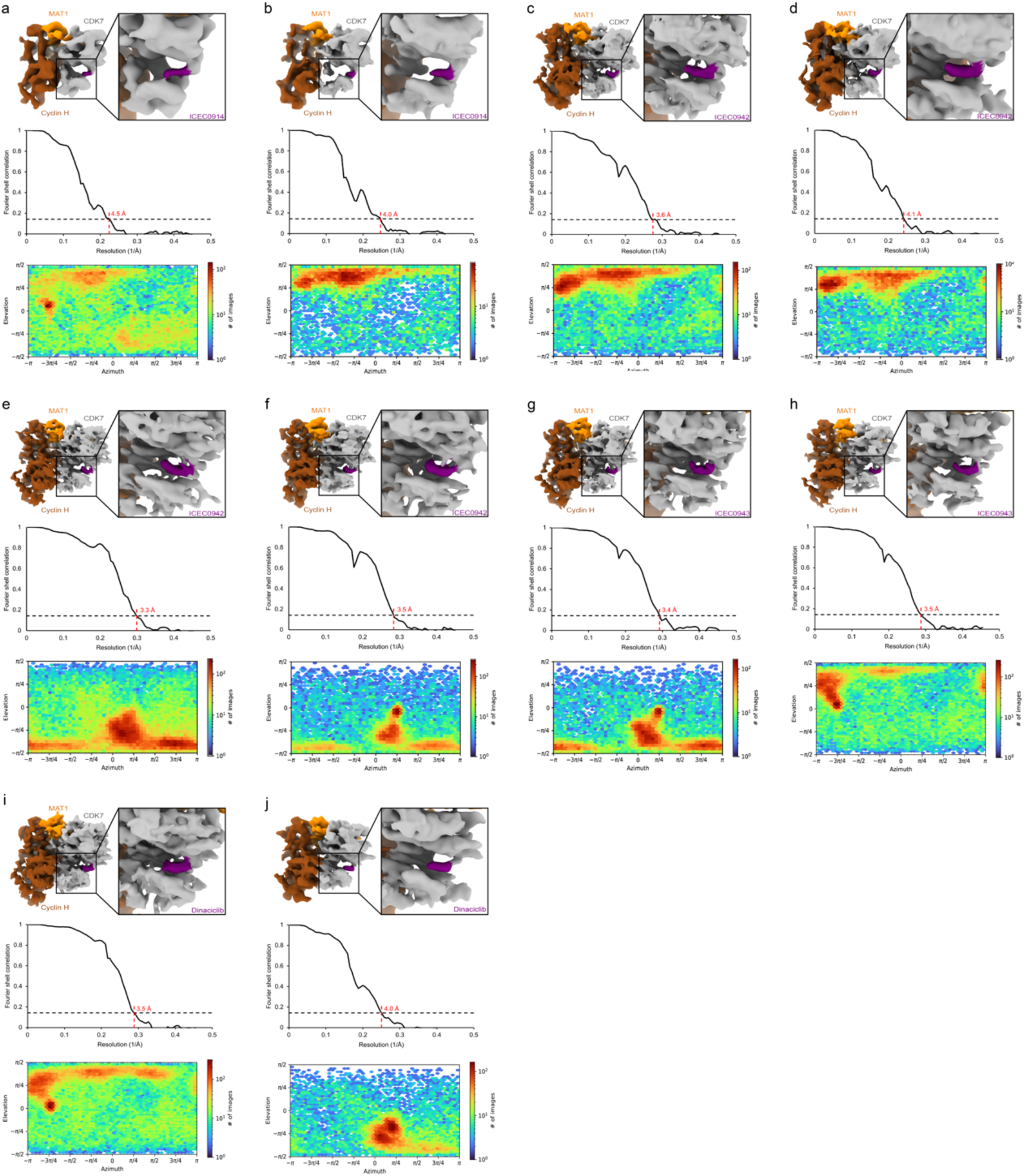
Live processing of 1-hour Glacios datasets, part 2. (**a**-**j**) Overview of 1-hour Glacios datasets. Top panels show a view of the 3D reconstruction (CDK7 grey, cyclin H brown, MAT1 orange) and a close-up view of the density for bound inhibitors (purple). Middle panels show the resolution according to the FSC = 0.143 threshold. Bottom panels show orientation distribution plots from cryoSPARC.

**Extended Data Figure 5.**
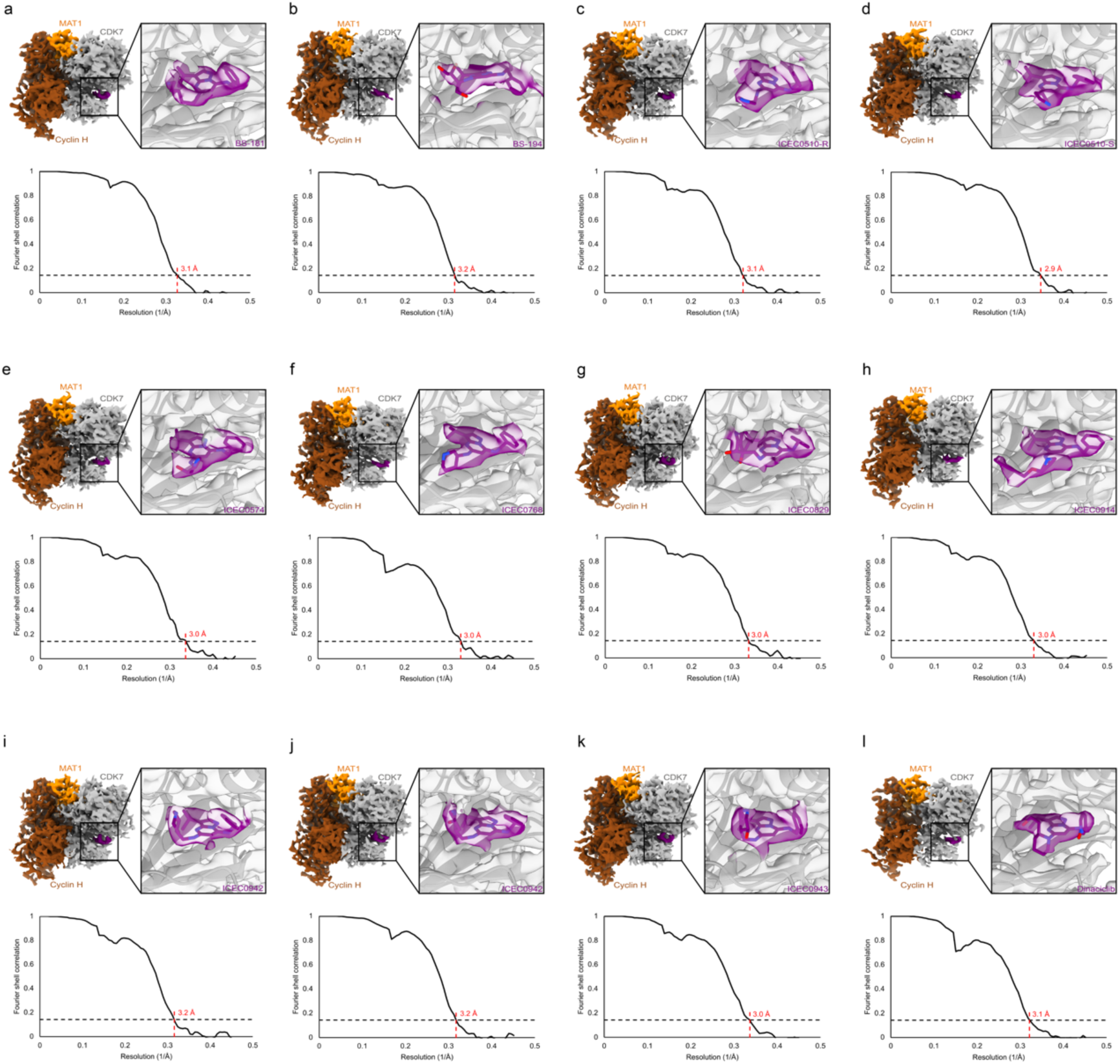
Processing of 4-hour Glacios datasets. (**a**-**l**) Overview of 4-hour Glacios dataset results. Top panels show a view of the 3D reconstruction (CDK7 grey, cyclin H brown, MAT1 orange) and a close-up view of the model and density for bound inhibitors (purple). Bottom panels show the resolution according to the FSC = 0.143 threshold.

**Extended Data Figure 6.**
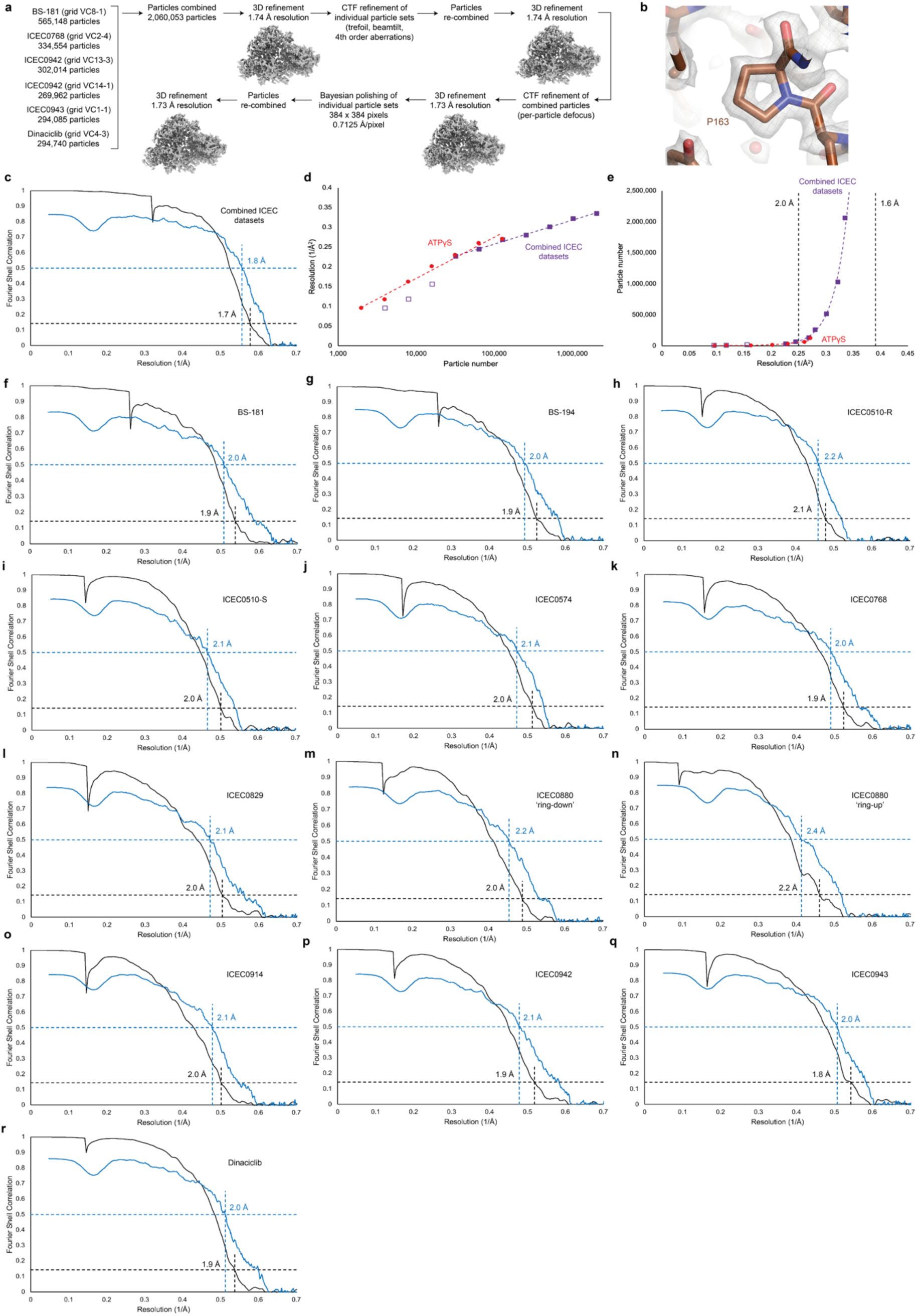
High-resolution cryo-EM structure of CAK and FSC curves. (**a**) Processing workflow leading to a 1.7 Å-reconstruction of the human CAK with averaged inhibitor density. (**b**) View of a proline residue (P163 of cyclin H) in the 1.7 Å-resolution reconstruction of the human CAK. (**c**) FSC curves for the 1.7 Å-reconstruction. The half-map FSC is shown in black, the model vs. map FSC is shown in blue. Resolution values are estimated from the FSC = 0.143 criterion for half-maps, and the FSC = 0.5 criterion for model vs. map FSCs ^39^. (**d**) Henderson-Rosenthal Plot for particle subsets derived by splitting the 1.7 Å- dataset in half 9 consecutive times (purple). At low particle numbers, the data deviate from linearity (outlined purple squares). These points were not used for extrapolation to higher particle numbers (see panel e). Additionally, data for the ATPγS dataset, which reached higher resolutions at low particle numbers, possibly due to the absence of DMSO in the sample buffer, is plotted in red. (**e**) The data from panel d plotted with particles on a linear scale to visualise the exponential increase in particle number required to reach higher resolution. (**f**- **r**) FSC curves for high-resolution inhibitor-bound structures.

**Extended Data Figure 7.**
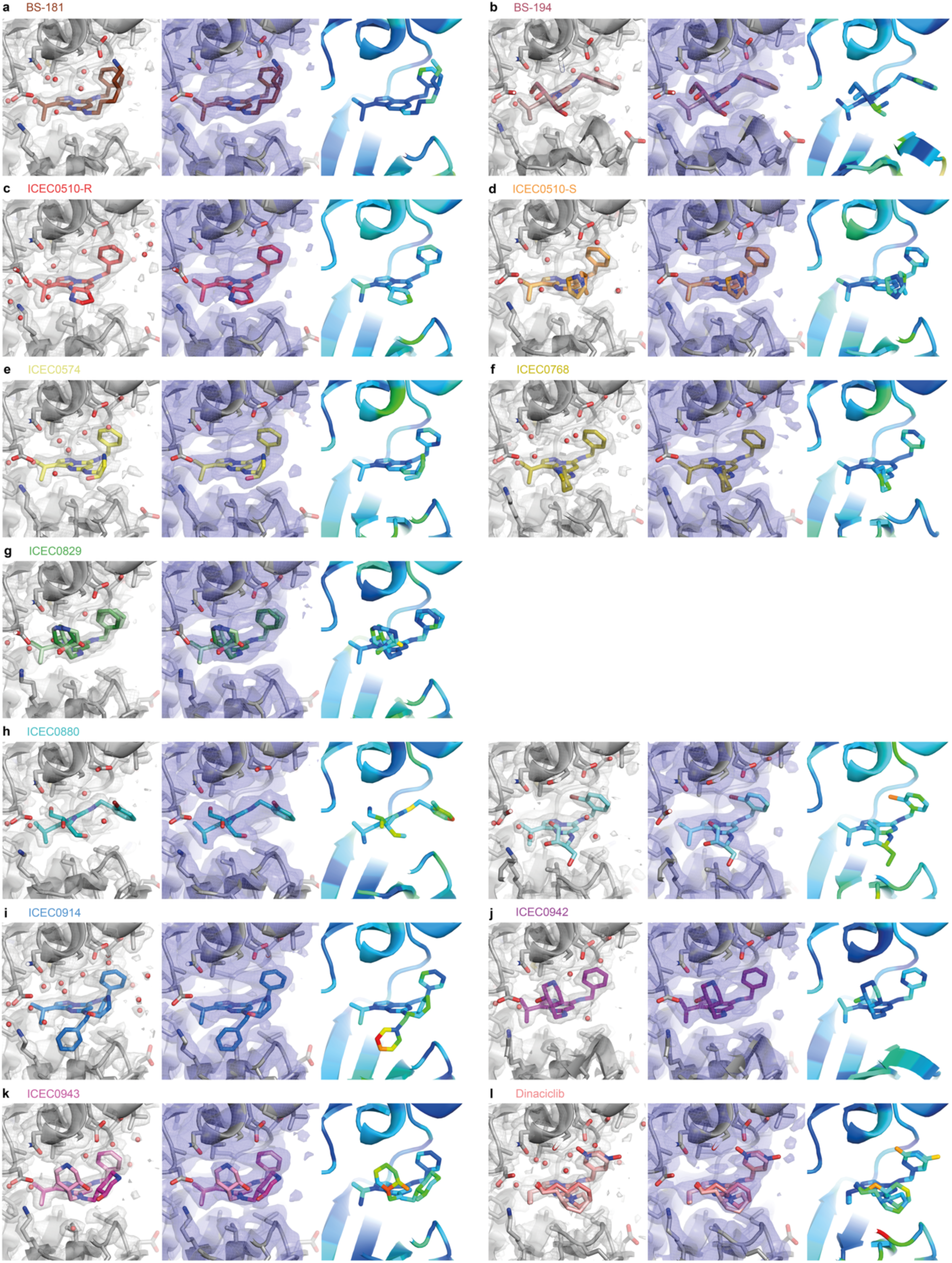
Structures of pyrazolopyrimidine-type inhibitors bound to CDK7 shown with the corresponding cryo-EM maps. (a) BS-181. (b) BS-194. (c) ICEC510-R. (d) ICEC510-S (two conformers). (e) ICEC0574. (f) ICEC0768. (g) ICEC0829 (two enantiomers). (h) ICEC0880 (two different positions correlated with conformational changes in CDK7). (i) ICEC0914. (**j**) The clinical inhibitor ICEC0942. (**k**) ICEC0943, an enantiomer of ICEC0942 (two conformers). (**l**) Dinaciclib (two conformers because the nitroxide position could not be assigned unambiguously). Panels show the fitted inhibitors in the post-processed cryo-EM map (grey, left side panels), inhibitors fitted in maps filtered to 3 Å resolution to visualise poorly ordered chemical groups (blue, middle panels), and inhibitors coloured by Q-scores (red to blue, right-hand panels).

**Extended Data Figure 8.**
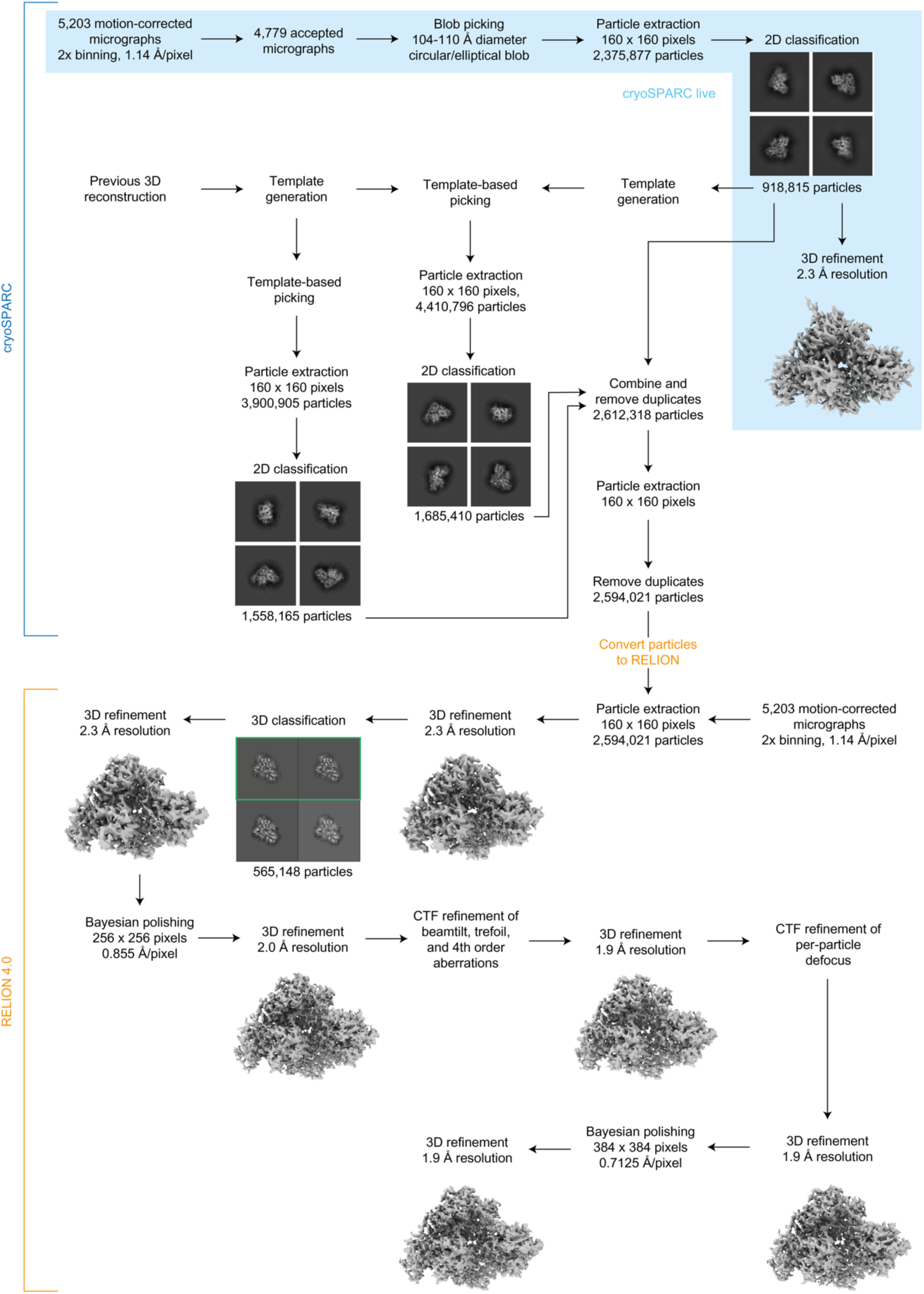
Data processing strategy. Processing stages using cryoSPARC and RELION are indicated. Steps performed in cryoSPARC live are shaded in blue.

**Extended Data Table 1.**
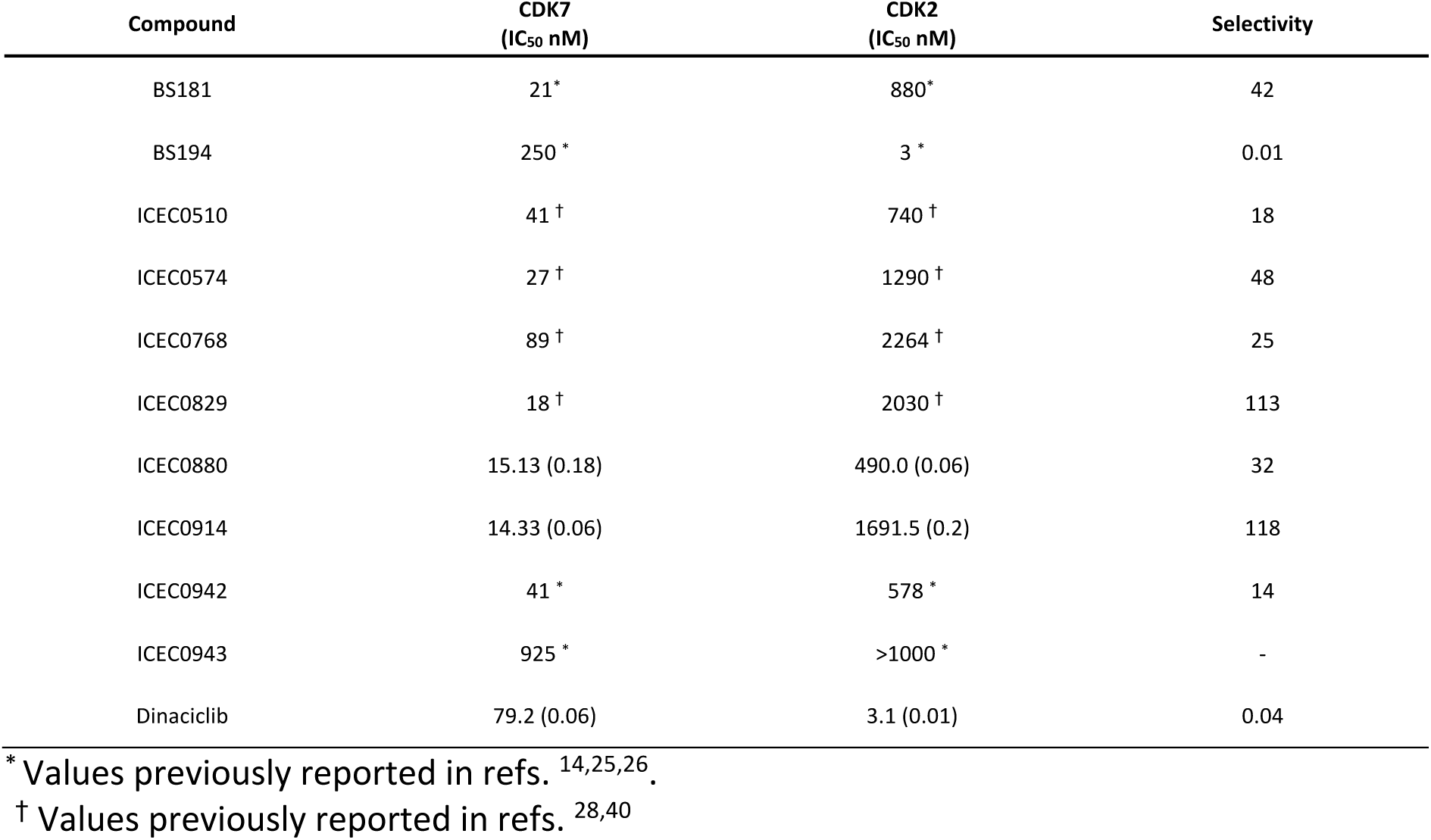
Characterisation of ICEC-series inhibitors. Half-maximal inhibitory concentrations (IC_50_) of pyrazolopyrimidine inhibitors used in this study, determined by *in vitro* kinase assays.

### Supplementary Information

**This file contains:**

Supplementary Notes 1-2

Supplementary Tables 1-10

### Supplementary Note 1 – Comparison of 200 kV Glacios 2 and 300 kV Titan Krios G4 instrument performance

While most high-resolution structures deposited to the Electron Microscopy Data Resource (http://www.emdataresource.org) have been determined from data collected on high-end, 300 kV electron microscopes, more accessible 200 kV microscopes equipped with autoloaders and direct electron detectors are increasingly applied to determination of published structures and atomic models. Their applicability to a variety of specimens, including small soluble complexes ^12, 41^ and small membrane proteins ^42^, and their ability to achieve better than 2 Å resolution on ideal – i.e. symmetrical, rigid, and often large – targets ^43, 44^ have been demonstrated. However, there is a scarcity of direct comparisons between the results obtained using 200 kV and 300 kV instruments on the exact same specimen.

To address this question and provide a more systematic comparison, we determined the structure of the CDK-activating kinase bound to an inhibitor (see main text) from 4,173 good micrographs (count after removal of poor-quality micrographs) collected on a 300 kV Titan Krios G4 cryo-TEM equipped with a cold-FEG, a Selectris X energy filter, and a Falcon 4i direct electron detector. Using a combination of cryoSPARC ^22^ and RELION ^23^ for image processing, we were able to reconstruct a 3D map at 2.0 Å from these data (Extended Data Fig. 2a, c). Having verified the ability to achieve high resolution from this specimen, we transferred the same grid to a 200 kV Glacios 2 cryo-TEM equipped with a Selectris X energy filter and a Falcon 4i direct electron detector and acquired 7,907 good micrographs (count after removal of poor-quality micrographs). The energy filter and detector were thus identical, while the electron source and the optics were different between these datasets. From this latter setup, we achieved 2.3 Å resolution (Extended Data Fig. 2a, b). This is an improvement compared to our previous best result at 2.5 Å using a 200 kV Talos Arctica cryo-TEM equipped with a Gatan K3 direct detector ^12^ and might be attributable to the use of a different camera and energy filtration. However, despite this improvement in the 200 kV-data, the result from the 300 kV- instrument was superior to the map derived from the 200 kV-data, notably from fewer micrographs (4,173 for the Titan Krios G4, 7,907 for the Glacios 2) and within a shorter collection time (10 hours and 22 hours, respectively). To facilitate analysis of these data we computed reconstructions from data subsets and plotted the resulting resolutions (Extended Data Fig. 2d, e).

We note that this result was obtained with a small (85 kDa) complex that lacks symmetry, a combination that is typically considered challenging ^16^, and is therefore more representative of high-end biological use cases than experiments with model protein complexes. Results with large or highly symmetric specimens, where alignment accuracy is less limiting due to greater signal, may show different resolution gaps between the two instrument types. We also acknowledge that this comparison is primarily valid for the specific system used, that the cold-FEG may contribute to the superior results from the 300 kV system, and that the DQE difference between 200 kV and 300 kV data may depend on the direct detector model used.

### Supplementary Note 2 – Comparison of the impact of energy filtration on the quality of low-defocus cryo-EM data of a small complex

The importance of energy filtration as a means to remove inelastically scattered electrons from the electron beam arriving at the detector, thereby improving the signal-to-noise ratio by reducing electrons that only contribute to the noise but not the signal, is well-established for thick specimens, such as those routinely found in cryo-electron tomography experiments ^45^. While many high-resolution single-particle structures obtained from thin specimens employed energy filtration as well ^20, 46^, there are few systematic tests that analyse the effect of energy filtration on the quality of data obtained from thin single-particle specimens of smaller complexes such as the CAK studied in this work. We therefore collected low-defocus (0.5-1.0 μm) data using the standard setup (Titan Krios G4 with cold-FEG, Selectris X energy filter, and Falcon 4i detector) and then proceeded to collect another dataset on the same grid after retraction of the energy filtration slit. We obtained a resolution of 2.0 Å from the energy filtered data and 2.3 Å from the data after retraction of the slit (Extended Data Fig. 2a). Data were processed as described for all other complexes (see Methods), except that additional instances of 2D classification were run to identify all suitable particles, and that only one round of Bayesian polishing was performed before the final refinement. Refinement of subsets showed that the energy filtered data also provide higher resolution for smaller dataset sizes, and that breaking the 2 Å barrier may be difficult without energy filtration for our specimen and data collection parameters (Extended Data Fig. 2f, g).

We note that our experiment strongly supports a positive effect of energy filtration on the data quality of the 85 kDa human CAK collected at low defocus, but that further experiments are required to investigate if the conclusions drawn from this comparison can be extrapolated to other specimens, such as complexes of larger size, and different data collection parameters, such as higher defocus.

**Supplementary Table 1.**
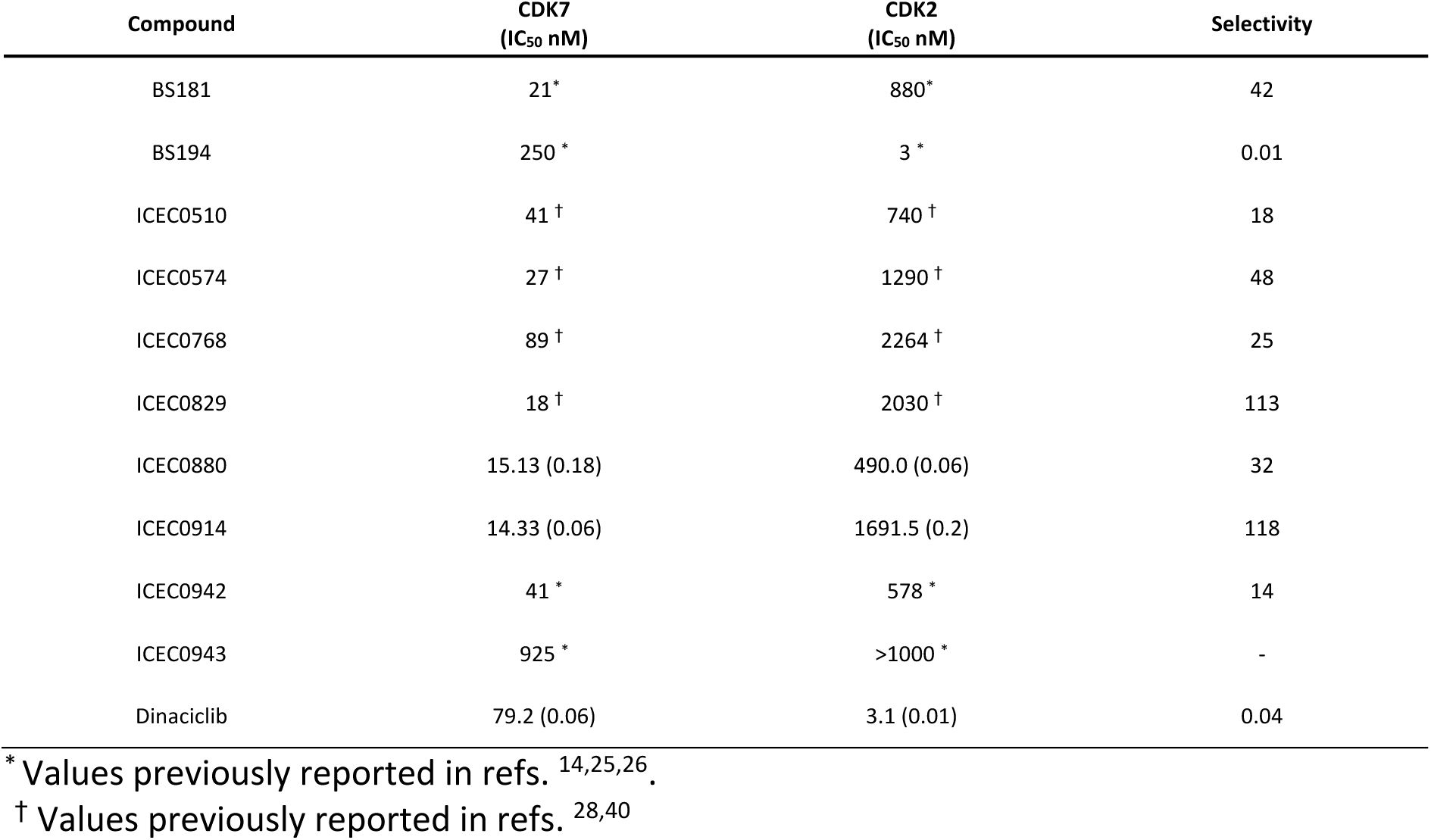
Glacios 2 1-hour screening datasets collected during this study. Grids selected for the next step in the data collection pipeline are marked in bold.

**Supplementary Table 2.** Glacios 2 4-hour screening datasets collected during this study.

| Ligand | Grid | Micrographs collected | Micrographs accepted | Particles refined (live) | Resolution cryoSPARC live (Å) | Particles refined (final) | Resolution RELION (Å) |
| --- | --- | --- | --- | --- | --- | --- | --- |
| BS-181 | VC8-2 | 1,686 | 1,281 | 201,719 | 3.0 | 93,194 | 3.1 |
| BS-194 | VC9-4 | 1,659 | 1,326 | 282,212 | 3.1 | 55,890 | 3.2 |
| ICEC0510-R | VC6-4 | 1,678 | 1,106 | 176,583 | 3.1 | 59,714 | 3.1 |
| ICEC0510-S | VC7-3 | 1,558 | 1,242 | 220,033 | 2.9 | 166,785 | 2.9 |
| ICEC0574 | VC11-4 | 1,764 | 953 | 156,086 | 3.0 | 62,893 | 3.0 |
| ICEC0768 | VC2-1 | 1,703 | 1,227 | 191,294 | 3.0 | 149,596 | 3.0 |
| ICEC0829 | VC12-3 | 1,594 | 1,110 | 144,402 | 3.0 | 60,827 | 3.0 |
| ICEC0880 | VC5-1* | 10,576 | 7,907 | 1,232,300 | 3.3 | 226,221 | 2.3 |
| ICEC0914 | VC3-1 | 1,632 | 1,217 | 145,713 | 3.0 | 47,422 | 3.0 |
| ICEC0942 | VC13-1 | 1,776 | 1,007 | 143,912 | 3.3 | 62,044 | 3.2 |
|  | VC14-1 | 1,658 | 1,285 | 246,409 | 3.1 | 134,727 | 3.2 |
| ICEC0943 | VC1-1 | 1,202 | 783 | 95,031 | 2.9 | 60,500 | 3.0 |
| Dinaciclib | VC4-1 | 1,637 | 1,194 | 160,000 | 3.2 | 128,172 | 3.1 |

\* full-size dataset for performance comparison with Titan Krios G4

**Supplementary Table 3.** Titan Krios G4 datasets collected during this study.

| Ligand | Grid | Micrographs collected | Micrographs accepted | Particles refined (live) | Resolution cryoSPARC (Å)* | Particles refined (final) | Resolution RELION (Å) |
| --- | --- | --- | --- | --- | --- | --- | --- |
| apo | VC15-1 | 5,805 | 5,036 | 1,402,951 | 2.3 | 486,457 | 2.3 |
| ATP<sub>γ</sub>S | VC16-3 | 5,781 | 5,238 | 830,780 | 2.3 | 126,177 | 1.9 |
| THZ1 | BG29-1 | 6,029 | 5,015 | 932,552 | 2.3 | 555,117 | 1.9 |
|  | BG29-2 | 6,542 | 5,553 | 850,654 | 2.3 |  |  |
| LDC4297 | BG30-3 | 9,553 | 6,426 | 835,340 | 2.5 | 273,097 | 2.1 |
| BS-181 | VC8-1 | 5,203 | 4,779 | 918,185 | 2.3 | 565,148 | 1.9 |
| BS-194 | VC9-4 | 6,351 | 5,725 | 1,041,685 | 2.3 | 688,511 | 1.9 |
| ICEC0510-R | VC6-4 | 6,309 | 5,130 | 981,580 | 2.5 | 268,647 | 2.1 |
| ICEC0510-S | VC7-3 | 5,081 | 3,564 | 916,457 | 2.3 | 233,209 | 2.0 |
| ICEC0574 | VC11-4 | 5,132 | 3,400 | 770,136 | 2.4 | 410,939 | 2.0 |
| ICEC0768 | VC2-4 | 5,583 | 4,038 | 803,921 | 2.3 | 334,554 | 1.9 |
| ICEC0829 | VC12-1 | 5,502 | 3,925 | 688,486 | 2.4 | 253,285 | 2.0 |
|  | VC12-3 | 5,079 | 1,990 | 390,255 | 2.5 | 171,002 | 2.0 |
|  | VC12-3<sup>‡</sup> | 5,167 | 2,457 | 211,647 | 2.8 | 165,849 | 2.4 |
| ICEC0880 | VC5-1 | 5,426 | 4,173 | 621,371 | 2.4 | 158,290 | 2.0 |
|  | State 1<sup>†</sup> |  |  |  |  | 112,809 | 2.0 |
|  | State 2<sup>†</sup> |  |  |  |  | 45,481 | 2.2 |
| ICEC0914 | VC3-1 | 5,202 | 3,823 | 709,706 | 2.3 | 237,743 | 2.0 |
| ICEC0942 | VC13-3 | 5,231 | 4,423 | 1,010,580 | 2.4 | 302,014 | 1.9 |
|  | VC14-1 | 5,367 | 3,885 | 790,254 | 2.3 | 269,962 | 1.9 |
| ICEC0943 | VC1-1 | 5,729 | 4,649 | 959,204 | 2.3 | 294,085 | 1.8 |
| Dinaciclib | VC4-3 | 5,210 | 4,561 | 776,307 | 2.3 | 294,740 | 1.9 |

\* limited to 2.28 Å (Nyquist frequency) in pre-processing due to 2x binning of input movies
<sup>†</sup> state 1: “ring-down” conformation; state 2: “ring-up” conformation
<sup>‡</sup> energy filter slit retracted

**Supplementary Table 4.**
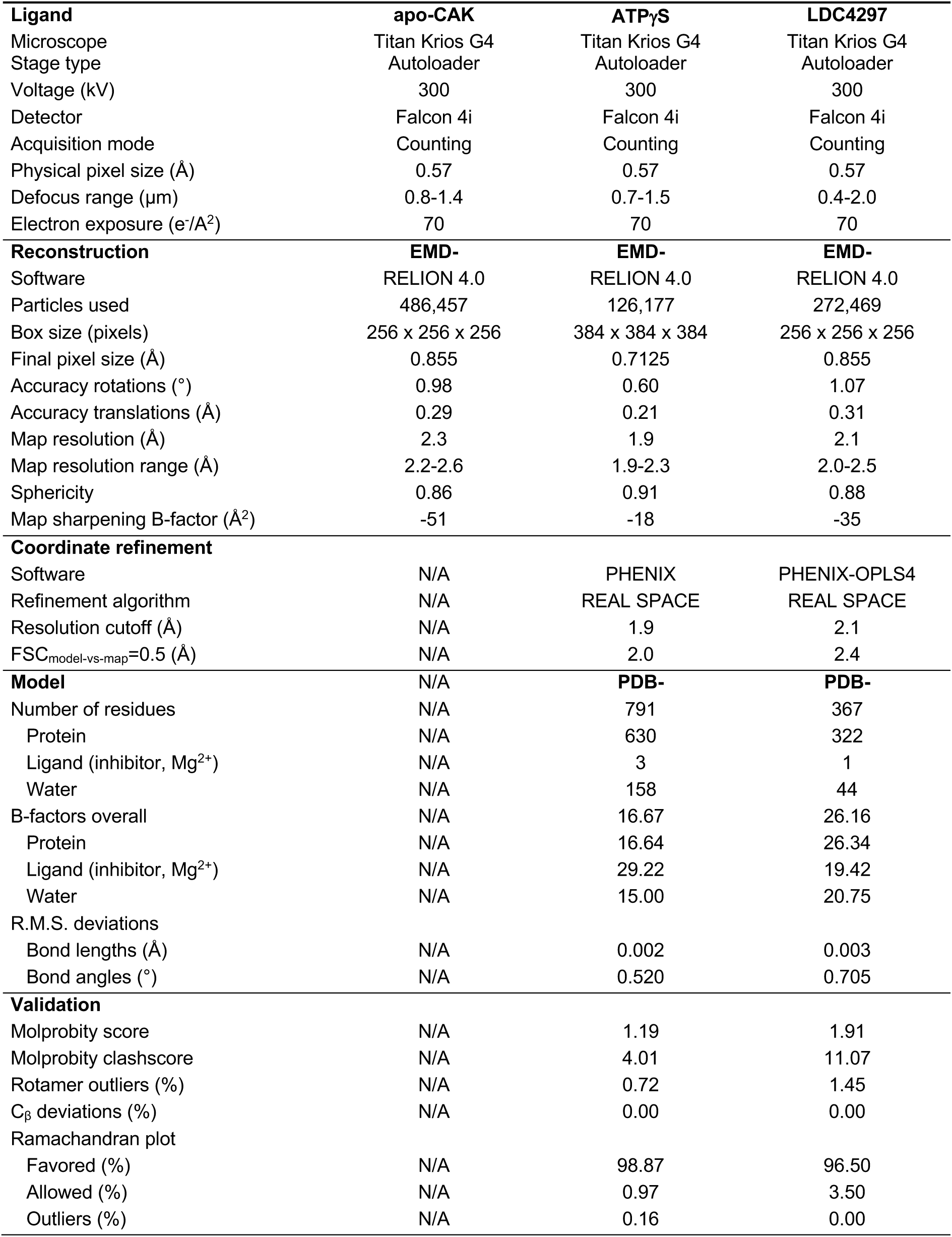
Cryo-EM data collection, 3D reconstruction, and refinement statistics, part 1.

| **Ligand** | **apo-CAK** | **ATPγS** | **LDC4297** |
| --- | --- | --- | --- |
| Microscope | Titan Krios G4 | Titan Krios G4 | Titan Krios G4 |
| Stage type | Autoloader | Autoloader | Autoloader |
| Voltage (kV) | 300 | 300 | 300 |
| Detector | Falcon 4i | Falcon 4i | Falcon 4i |
| Acquisition mode | Counting | Counting | Counting |
| Physical pixel size (Å) | 0.57 | 0.57 | 0.57 |
| Defocus range (μm) | 0.8-1.4 | 0.7-1.5 | 0.4-2.0 |
| Electron exposure (e<sup>-</sup>/Å<sup>2</sup>) | 70 | 70 | 70 |
| **Reconstruction** | **EMD-** | **EMD-** | **EMD-** |
| Software | RELION 4.0 | RELION 4.0 | RELION 4.0 |
| Particles used | 486,457 | 126,177 | 272,469 |
| Box size (pixels) | 256 x 256 x 256 | 384 x 384 x 384 | 256 x 256 x 256 |
| Final pixel size (Å) | 0.855 | 0.7125 | 0.855 |
| Accuracy rotations (°) | 0.98 | 0.60 | 1.07 |
| Accuracy translations (Å) | 0.29 | 0.21 | 0.31 |
| Map resolution (Å) | 2.3 | 1.9 | 2.1 |
| Map resolution range (Å) | 2.2-2.6 | 1.9-2.3 | 2.0-2.5 |
| Sphericity | 0.86 | 0.91 | 0.88 |
| Map sharpening B-factor (Å<sup>2</sup>) | -51 | -18 | -35 |
| **Coordinate refinement** |  |  |  |
| Software | N/A | PHENIX | PHENIX-OPLS4 |
| Refinement algorithm | N/A | REAL SPACE | REAL SPACE |
| Resolution cutoff (Å) | N/A | 1.9 | 2.1 |
| FSC<sub>model-vs-map</sub>=0.5 (Å) | N/A | 2.0 | 2.4 |
| **Model** | **N/A** | **PDB-** | **PDB-** |
| Number of residues | N/A | 791 | 367 |
| Protein | N/A | 630 | 322 |
| Ligand (inhibitor, Mg<sup>2+</sup>) | N/A | 3 | 1 |
| Water | N/A | 158 | 44 |
| B-factors overall | N/A | 16.67 | 26.16 |
| Protein | N/A | 16.64 | 26.34 |
| Ligand (inhibitor, Mg<sup>2+</sup>) | N/A | 29.22 | 19.42 |
| Water | N/A | 15.00 | 20.75 |
| R.M.S. deviations |  |  |  |
| Bond lengths (Å) | N/A | 0.002 | 0.003 |
| Bond angles (°) | N/A | 0.520 | 0.705 |
| **Validation** |  |  |  |
| Molprobability score | N/A | 1.19 | 1.91 |
| Molprobability clashscore | N/A | 4.01 | 11.07 |
| Rotamer outliers (%) | N/A | 0.72 | 1.45 |
| Cβ deviations (%) | N/A | 0.00 | 0.00 |
| Ramachandran plot |  |  |  |
| Favored (%) | N/A | 98.87 | 96.50 |
| Allowed (%) | N/A | 0.97 | 3.50 |
| Outliers (%) | N/A | 0.16 | 0.00 |

**Supplementary Table 5.**
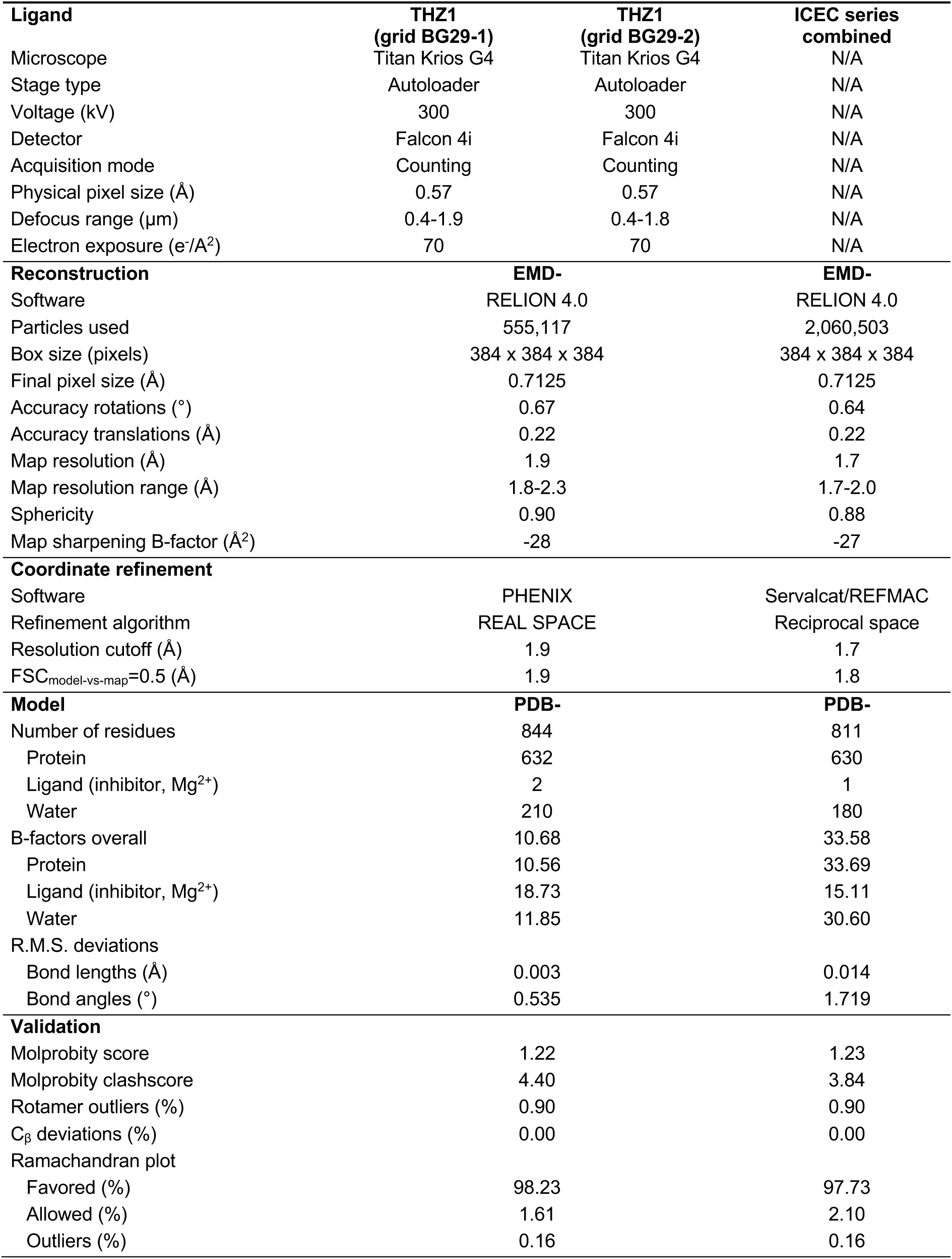
Cryo-EM data collection, 3D reconstruction, and refinement statistics, part 2.

**Supplementary Table 6.** Cryo-EM data collection, 3D reconstruction, and refinement statistics, part 3.

| **Ligand** | **BS-181** | **BS-194** | **ICEC0510-R** |
| --- | --- | --- | --- |
| Microscope | Titan Krios G4 | Titan Krios G4 | Titan Krios G4 |
| Stage type | Autoloader | Autoloader | Autoloader |
| Voltage (kV) | 300 | 300 | 300 |
| Detector | Falcon 4i | Falcon 4i | Falcon 4i |
| Acquisition mode | Counting | Counting | Counting |
| Physical pixel size (Å) | 0.57 | 0.57 | 0.57 |
| Defocus range (µm) | 0.4-1.5 | 0.4-1.8 | 0.4-1.6 |
| Electron exposure (e<sup>-</sup>/Å<sup>2</sup>) | 70 | 70 | 70 |
| **Reconstruction** | **EMD-** | **EMD-** | **EMD-** |
| Software | RELION 4.0 | RELION 4.0 | RELION 4.0 |
| Particles used | 565,148 | 688,511 | 268,647 |
| Box size (pixels) | 384 x 384 x 384 | 384 x 384 x 384 | 384 x 384 x 384 |
| Final pixel size (Å) | 0.7125 | 0.7125 | 0.7125 |
| Accuracy rotations (°) | 0.68 | 0.84 | 0.71 |
| Accuracy translations (Å) | 0.23 | 0.25 | 0.23 |
| Map resolution (Å) | 1.9 | 1.9 | 2.1 |
| Map resolution range | 1.8-2.3 | 1.8-2.4 | 2.0-2.5 |
| Sphericity | 0.84 | 0.86 | 0.86 |
| Map sharpening B-factor (Å<sup>2</sup>) | -27 | -26 | -28 |
| **Coordinate refinement** |  |  |  |
| Software | PHENIX | PHENIX | PHENIX |
| Refinement algorithm | REAL SPACE | REAL SPACE | REAL SPACE |
| Resolution cutoff (Å) | 1.9 | 1.9 | 2.1 |
| FSC<sub>model-vs-map</sub>=0.5 (Å) | 2.0 | 2.0 | 2.2 |
| **Model** | **PDB-** | **PDB-** | **PDB-** |
| Number of residues | 771 | 778 | 760 |
| Protein | 630 | 631 | 630 |
| Ligand (inhibitor, Mg<sup>2+</sup>) | 2 | 2 | 2 |
| Water | 139 | 145 | 128 |
| B-factors overall | 7.88 | 23.46 | 22.22 |
| Protein | 7.93 | 23.53 | 22.29 |
| Ligand (inhibitor, Mg<sup>2+</sup>) | 5.55 | 32.16 | 19.14 |
| Water | 6.27 | 17.63 | 20.04 |
| R.M.S. deviations |  |  |  |
| Bond lengths (Å) | 0.004 | 0.003 | 0.004 |
| Bond angles (°) | 0.639 | 0.570 | 0.617 |
| **Validation** |  |  |  |
| Molprobrity score | 1.45 | 1.45 | 1.48 |
| Molprobrity clashscore | 7.06 | 7.81 | 7.36 |
| Rotamer outliers (%) | 0.72 | 0.54 | 1.26 |
| C<sub>β</sub> deviations (%) | 0.00 | 0.00 | 0.00 |
| Ramachandran plot |  |  |  |
| Favored (%) | 97.73 | 97.90 | 98.38 |
| Allowed (%) | 2.10 | 1.94 | 1.46 |
| Outliers (%) | 0.16 | 0.16 | 0.16 |

**Supplementary Table 7.** Cryo-EM data collection, 3D reconstruction, and refinement statistics, part 4.

| **Ligand** | **ICEC0510-S** | **ICEC0574** | **ICEC0768** |
| --- | --- | --- | --- |
| Microscope | Titan Krios G4 | Titan Krios G4 | Titan Krios G4 |
| Stage type | Autoloader | Autoloader | Autoloader |
| Voltage (kV) | 300 | 300 | 300 |
| Detector | Falcon 4i | Falcon 4i | Falcon 4i |
| Acquisition mode | Counting | Counting | Counting |
| Physical pixel size (Å) | 0.57 | 0.57 | 0.57 |
| Defocus range (µm) | 0.4-1.9 | 0.4-1.8 | 0.5-1.7 |
| Electron exposure (e<sup>-</sup>/Å<sup>2</sup>) | 70 | 70 | 70 |
| **Reconstruction** | **EMD-** | **EMD-** | **EMD-** |
| Software | RELION 4.0 | RELION 4.0 | RELION 4.0 |
| Particles used | 233,209 | 410,939 | 334,554 |
| Box size (pixels) | 384 x 384 x 384 | 384 x 384 x 384 | 384 x 384 x 384 |
| Final pixel size (Å) | 0.7125 | 0.7125 | 0.7125 |
| Accuracy rotations (°) | 0.67 | 0.75 | 0.68 |
| Accuracy translations (Å) | 0.22 | 0.25 | 0.23 |
| Map resolution (Å) | 2.0 | 2.0 | 1.9 |
| Map resolution range | 2.0-2.2 | 1.9-2.7 | 1.9-2.7 |
| Sphericity | 0.92 | 0.84 | 0.84 |
| Map sharpening B-factor (Å<sup>2</sup>) | -21 | -25 | -24 |
| **Coordinate refinement** |  |  |  |
| Software | PHENIX | PHENIX | PHENIX |
| Refinement algorithm | REAL SPACE | REAL SPACE | REAL SPACE |
| Resolution cutoff (Å) | 2.0 | 2.0 | 1.9 |
| FSC<sub>model-vs-map</sub>=0.5 (Å) | 2.1 | 2.1 | 2.0 |
| **Model** | **PDB-** | **PDB-** | **PDB-** |
| Number of residues | 786 | 747 | 757 |
| Protein | 630 | 630 | 630 |
| Ligand (inhibitor, Mg<sup>2+</sup>) | 2 | 2 | 2 |
| Water | 154 | 115 | 125 |
| B-factors overall | 29.55 | 20.71 | 16.52 |
| Protein | 29.81 | 20.85 | 16.59 |
| Ligand (inhibitor, Mg<sup>2+</sup>) | 19.23 | 13.49 | 15.59 |
| Water | 24.77 | 16.40 | 13.67 |
| R.M.S. deviations |  |  |  |
| Bond lengths (Å) | 0.003 | 0.003 | 0.003 |
| Bond angles (°) | 0.527 | 0.520 | 0.623 |
| **Validation** |  |  |  |
| Molprobtity score | 1.36 | 1.41 | 1.46 |
| Molprobtity clashscore | 6.55 | 7.55 | 8.63 |
| Rotamer outliers (%) | 0.72 | 0.90 | 0.90 |
| C<sub>β</sub> deviations (%) | 0.00 | 0.00 | 0.00 |
| Ramachandran plot |  |  |  |
| Favored (%) | 98.54 | 98.38 | 98.06 |
| Allowed (%) | 1.29 | 1.46 | 1.78 |
| Outliers (%) | 0.16 | 0.16 | 0.16 |

**Supplementary Table 8.**
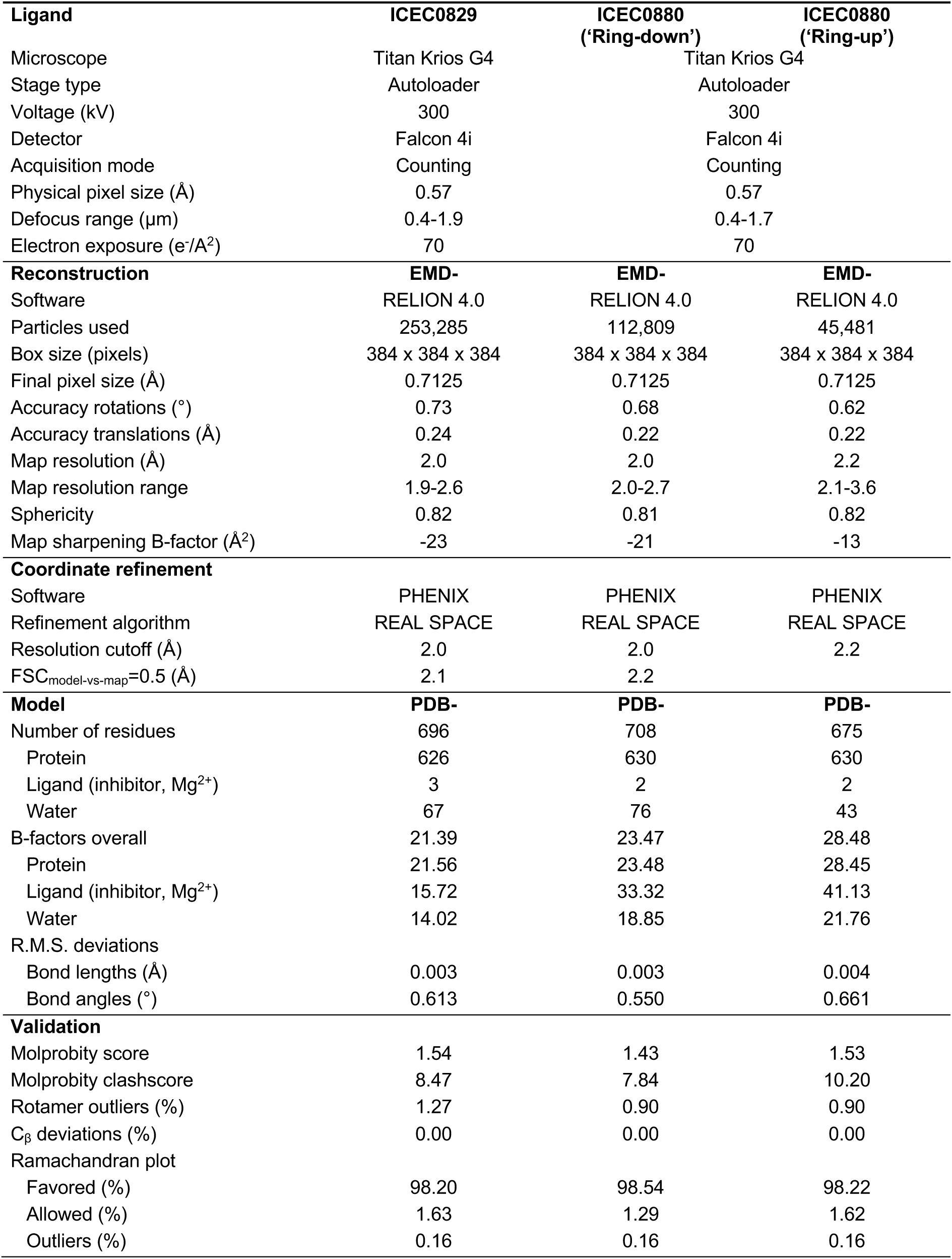
Cryo-EM data collection, 3D reconstruction, and refinement statistics, part 5.

**Supplementary Table 9.** Cryo-EM data collection, 3D reconstruction, and refinement statistics, part 6.

| **Ligand** | **ICEC0914** | **ICEC0942** | **ICEC0943** |
| --- | --- | --- | --- |
| Microscope | Titan Krios G4 | Titan Krios G4 | Titan Krios G4 |
| Stage type | Autoloader | Autoloader | Autoloader |
| Voltage (kV) | 300 | 300 | 300 |
| Detector | Falcon 4i | Falcon 4i | Falcon 4i |
| Acquisition mode | Counting | Counting | Counting |
| Physical pixel size (Å) | 0.57 | 0.57 | 0.57 |
| Defocus range (µm) | 0.4-2.0 | 0.4-1.9 | 0.4-1.7 |
| Electron exposure (e<sup>-</sup>/Å<sup>2</sup>) | 70 | 70 | 70 |
| **Reconstruction** | **EMD-** | **EMD-** | **EMD-** |
| Software | RELION 4.0 | RELION 4.0 | RELION 4.0 |
| Particles used | 237,743 | 269,962 | 294,085 |
| Box size (pixels) | 384 x 384 x 384 | 384 x 384 x 384 | 384 x 384 x 384 |
| Final pixel size (Å) | 0.7125 | 0.7125 | 0.7125 |
| Accuracy rotations (°) | 0.73 | 0.69 | 0.67 |
| Accuracy translations (Å) | 0.24 | 0.22 | 0.22 |
| Map resolution (Å) | 2.0 | 1.9 | 1.8 |
| Map resolution range | 1.9-2.5 | 1.9-2.5 | 1.8-2.4 |
| Sphericity | 0.83 | 0.84 | 0.85 |
| Map sharpening B-factor (Å<sup>2</sup>) | -24 | -23 | -19 |
| **Coordinate refinement** |  |  |  |
| Software | PHENIX | PHENIX, OPLS4 | PHENIX |
| Refinement algorithm | REAL SPACE | REAL SPACE | REAL SPACE |
| Resolution cutoff (Å) | 2.0 | 1.9 | 1.8 |
| FSC<sub>model-vs-map</sub>=0.5 (Å) | 2.1 | 2.1 | 2.0 |
| **Model** | **PDB-** | **PDB-** | **PDB-** |
| Number of residues | 784 | 770 | 769 |
| Protein | 630 | 630 | 630 |
| Ligand (inhibitor, Mg<sup>2+</sup>) | 2 | 2 | 2 |
| Water | 152 | 138 | 137 |
| B-factors overall | 18.82 | 21.42 | 21.32 |
| Protein | 18.81 | 21.52 | 21.49 |
| Ligand (inhibitor, Mg<sup>2+</sup>) | 19.44 | 14.52 | 15.53 |
| Water | 18.87 | 19.22 | 17.77 |
| R.M.S. deviations |  |  |  |
| Bond lengths (Å) | 0.004 | 0.003 | 0.003 |
| Bond angles (°) | 0.620 | 0.640 | 0.662 |
| **Validation** |  |  |  |
| Molprobrity score | 1.40 | 1.48 | 1.36 |
| Molprobrity clashscore | 6.86 | 6.57 | 6.45 |
| Rotamer outliers (%) | 1.08 | 1.44 | 0.90 |
| C<sub>β</sub> deviations (%) | 0.00 | 0.00 | 0.00 |
| Ramachandran plot |  |  |  |
| Favored (%) | 98.38 | 98.22 | 98.38 |
| Allowed (%) | 1.46 | 1.62 | 1.46 |
| Outliers (%) | 0.16 | 0.16 | 0.16 |

**Supplementary Table 10.** Cryo-EM data collection, 3D reconstruction, and refinement statistics, part 7.

| **Ligand** | **Dinaciclib** |
| --- | --- |
| Microscope | Titan Krios G4 |
| Stage type | Autoloader |
| Voltage (kV) | 300 |
| Detector | Falcon 4i |
| Acquisition mode | Counting |
| Physical pixel size (Å) | 0.57 |
| Defocus range (µm) | 0.4-1.8 |
| Electron exposure (e<sup>-</sup>/Å<sup>2</sup>) | 70 |
| **Reconstruction** | **EMD-** |
| Software | RELION 4.0 |
| Particles used | 294,740 |
| Box size (pixels) | 384 x 384 x 384 |
| Final pixel size (Å) | 0.7125 |
| Accuracy rotations (°) | 0.60 |
| Accuracy translations (Å) | 0.21 |
| Map resolution (Å) | 1.9 |
| Map resolution range | 1.8-2.1 |
| Sphericity | 0.91 |
| Map sharpening B-factor (Å<sup>2</sup>) | -21 |
| **Coordinate refinement** |  |
| Software | PHENIX |
| Refinement algorithm | REAL SPACE |
| Resolution cutoff (Å) | 1.9 |
| FSC<sub>model-vs-map</sub>=0.5 (Å) | 2.0 |
| **Model** | **PDB-** |
| Number of residues | 804 |
| Protein | 630 |
| Ligand (inhibitor, Mg<sup>2+</sup>) | 2 |
| Water | 172 |
| B-factors overall | 15.14 |
| Protein | 15.12 |
| Ligand (inhibitor, Mg<sup>2+</sup>) | 16.40 |
| Water | 15.36 |
| R.M.S. deviations |  |
| Bond lengths (Å) | 0.004 |
| Bond angles (°) | 0.652 |
| **Validation** |  |
| Molprobit score | 1.38 |
| Molprobit clashscore | 6.84 |
| Rotamer outliers (%) | 0.54 |
| C<sub>β</sub> deviations (%) | 0.00 |
| Ramachandran plot |  |
| Favored (%) | 98.06 |
| Allowed (%) | 1.78 |
| Outliers (%) | 0.16 |

## Notes

### Summary of Updates

Errors in the author list and acknowledgments have been corrected.

